# Single-cell and Spatial Transcriptomics Clustering with an Optimized Adaptive K-Nearest Neighbor Graph

**DOI:** 10.1101/2023.10.13.562261

**Authors:** Jia Li, Yu Shyr, Qi Liu

**Affiliations:** Department of Biostatistics, Vanderbilt University Medical Center, Nashville, TN, 37203, USA; Center for Quantitative Sciences, Vanderbilt University Medical Center, Nashville, TN, 37232, USA

## Abstract

Single-cell and spatial transcriptomics have been widely used to characterize cellular landscape in complex tissues. To understand cellular heterogeneity, one essential step is to define cell types through unsupervised clustering. While typical clustering methods have difficulty in identifying rare cell types, approaches specifically tailored to detect rare cell types gain their ability at the cost of poorer performance for grouping abundant ones. Here, we developed aKNNO, a method to identify abundant and rare cell types simultaneously based on an adaptive k-nearest neighbor graph with optimization. Benchmarked on 38 simulated and 20 single-cell and spatial transcriptomics datasets, aKNNO identified both abundant and rare cell types accurately. Without sacrificing performance for clustering abundant cell types, aKNNO discovered known and novel rare cell types that those typical and even specifically tailored methods failed to detect. aKNNO, using transcriptome alone, stereotyped fine-grained anatomical structures more precisely than those integrative approaches combining expression with spatial locations and histology image.

## Introduction

Single-cell and spatial transcriptomics provide an unprecedented opportunity to navigate cellular landscape in complex tissues^1-4^. To understand cellular heterogeneity, one essential step is to define cell types through unsupervised clustering based on transcriptome similarity^5^. A number of clustering methods have been developed, most of which are generic algorithms adapted for single-cell transcriptomics analysis, such as k-means, hierarchical, density-based^6^, and community-detection-based clustering^7^. For example, RaceID^8^, SIMLR^9^, and SC3^10^ refine k-means for robust cell clustering. CIDR^11^, BackSPIN^12^, and pcaReduce^13^ extend hierarchical clustering to improve grouping ability on single cell transcriptomics. Phenograph^14^, Seurat^15^, and scanpy^16^ apply community-detection methods to define cell clusters. Although those typical methods achieved good performance in identifying abundant cell types, they all face the challenge of detecting rare ones.

To address the challenge, several approaches have been specifically designed or tailored to detect rare cell types, such as RaceID^8^, GiniClust^17^, GiniClust3^18^, FiRE^19^, and GapClust^20^. RaceID supplements k-means clustering with outlier detection to identify rare cell types^8^. GiniClust selects genes with high Gini index and then discovers rare cells based on density-based clustering^17^. GiniClust3 extends the method to identify both abundant and rare cell clusters using a cluster-aware, weighted ensemble approach^18^. FiRE uses the Sketching technique to assign a rareness score to each cell^19^. GapClust captures the abrupt local distances change to find rare cell clusters^20^. Those methods either only target rare cells but ignore abundant ones, or gain the ability of identifying rare cells at the cost of poorer performance for clustering abundant ones^21^.

Here, we propose aKNNO, which builds an optimized adaptive *k*-nearest neighbor graph for community-based detection. Compared to the traditional *k*-nearest neighbor (*k*NN) graph requiring a prespecified and fixed *k* for all cells, aKNNO chooses *k* adaptively for each cell based on its local distance distribution. The adaptive strategy enables the accurate detection of both abundant and rare cell types in a single run. The optimization step gets the most of the adaptive strategy and further improves rare cells identification. Benchmarked on 38 simulated and 17 single-cell transcriptomics datasets, aKNNO outperformed other methods for rare cells identification without scarifying the performance on abundant cells clustering. Applied on three spatial transcriptomics datasets, aKNNO stereotyped fine-grained anatomical patterns using gene expression alone, some of which were even missed by those methods integrating gene expression, spatial locations, and histology image.

## Results

### Overview of aKNNO

The community-detection methods have become increasingly popular, particularly for analyzing large single-cell transcriptomics datasets^21^. They first construct a *k*NN graph by connecting each cell to its nearest *k* cells measured by transcriptome similarity, and then group cells with dense connection. The choice of *k* has great impact on the clustering performance. A large *k* may generate phony connections between rare cells and cells in other clusters, while a small *k* may lead to overclustering of abundant cell types due to dominance of local variances. Cell populations in single-cell data are generally highly imbalanced, including both abundant and rare cells. Therefore, the traditional *k*NN graph using a universal *k* for all cells is unable to capture the inherent cellular structure accurately.

Instead of using a single *k* value for all cells, aKNNO chooses *k* adaptively for each cell based on its local distance. It automatically assigns a small *k* for rare cells to remove spurious long-range connections and connect only true nearest neighbors, and a large *k* for abundant cells to balance local and global variances. We used a toy example to illustrate how *k* is chosen adaptively (Fig. 1). In the example, K*max* is set to 10, meaning that the choice of *k* ranges from 1 to 10. For each cell, aKNNO finds its 10-nearest neighbors and sorts the distance in an ascending order (d1<d2<…<d10). *k*=K*max*=10 if d10<dcutoff; otherwise, *k* is chosen if d*k*<dcutoff and d*k+1*≥dcutoff. dcutoff is determined by 10-nearest distances of the cell and tuned by a hyperparameter *σ* (dcutoff=f(d1,..d10,*σ*)). As an example, cell A comes from a rare cluster containing only six cells, thus its 10-nearest distances increase dramatically from d5 to d6, leading to its 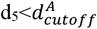 and 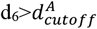. Therefore, *k* of 5 is chosen for the cell A (*k*A=5). As another example, cell B belongs to an abundant cell type, where its 10-nearest distances have a slow increase from d1 to d10, resulting in its 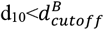. In this case, *k* of 10 is selected for the cell B (*k*B=10). In this way, aKNNO assigns the adaptive and optimal neighbors for each cell. To improve the robustness, the adaptive nearest-neighbor graph is reweighted based on the shared nearest neighbors of pairs of cells (SNN). Finally, Louvain community detection method is applied on the shared nearest neighbor graph to identify clusters (Fig. 1). The hyperparameter *σ* controls the sensitivity of aKNNO to the local distance change. aKNNO performs a grid search to find the optimal *σ* that balances the sensitivity and specificity of rare cluster identification (Fig. 1).

**Figure 1.**
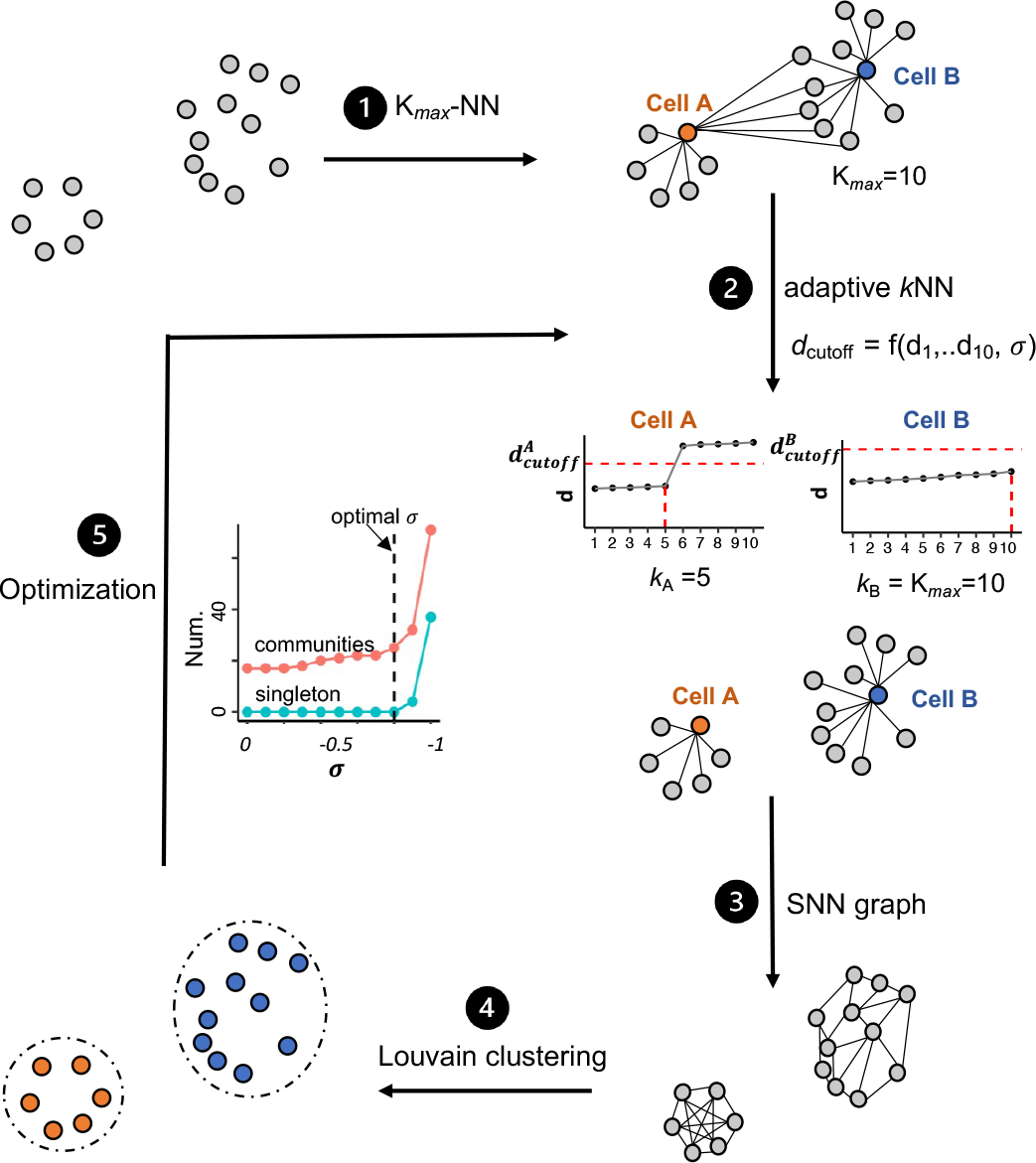
Overview of aKNNO. aKNNO includes five steps: 1) calculating the K*max*-nearest neighbors for all cells; 2) choosing *k* adaptively for each cell based on its local distance distribution; 3) building the shared nearest neighbor graph; 4) Clustering based on Louvain community-detection; 5) optimizing the hyperparameter *σ* by grid search.

### Performance of aKNNO in simulated datasets

We first compared the performance of aKNNO to the traditional *k*NN-based method in the Seurat^15^ (denoted as KNN) using simulated datasets with ground truth cell-type identity. To make a fair comparison, aKNNO and KNN processed the data exactly the same way except that aKNNO clustered cells based on an adaptive *k*-nearest neighbor graph while KNN grouped cells from a *k*NN graph with a fix *k* for all cells (default *k*=20, details in methods). We generated simulated datasets using a public single-cell RNA-seq dataset with approximately 68,000 peripheral blood mononuclear cells (PBMC68k)^22^ and 11 cell types. We simulated two settings, each of which contained two abundant and one rare cell types. In one setting, the two abundant cell types were CD19+ B cells (n=300) and CD56+ NK (n=200), while the rare cell type was CD8+ Cytotoxic T cells. In the other setting, the two abundant cell types were CD8+ Cytotoxic T cells (n=300) and CD56+ NK (n=200), while the rare cell type was CD19+ B cells. The first setting was more challenging than the second since its rare cells (CD8+ Cytotoxic T) were similar to one of abundant cell types (CD56+ NK), indicating that those rare cells were more likely to be hidden by abundant ones. We simulated 19 scenarios for each setting with the number of rare cells ranging from 2 to 20, and we generated 50 datasets for each scenario by random sampling cells from the PBMC68k.

We used F1-score for performance evaluation on rare cells identification, which reflects the balances between precision and sensitivity. The clustering results were obtained at five different resolutions (r=0.01, 0.8, 1, 1.5, and 2). aKNNO detected rare cells perfectly in all datasets with F1 scores of 1, no matter which resolution was used (Figs. 2a and 2b). It also grouped abundant cell types accurately and achieved the perfect agreement with the ground truth (adjusted rand index ARI=1) (Supplementary Fig. S1). In comparison, KNN completely failed to detect rare cells at every resolution when the number of rare cells was less than 11 in the first challenging setting (Fig. 2a). An example with 10 rare CD8+ Cytotoxic T cells was given in the Fig. 2c. Even at the lowest resolution of 0.01, aKNNO identified three clusters with 100% accuracy. KNN, however, grouped the 10 CD8+ cytotoxic T cells into CD56+ NK cells even at the highest resolution (KNN_high in the Fig. 2c). Moreover, the performance of KNN was sensitive to the resolution. For example, KNN achieved a F1 score of 1 at the resolution of 2 when there were 17 rare cells, while the F1 score was less than 0.05 at the resolution of 0.01. However, the high F1 score at the high resolution was gained at the cost of extremely overclustering of abundant cell types (KNN_high in the Supplementary Fig. S2). In the second setting, KNN failed to detect rare cells when the number of rare cells was less than nine (Fig. 2b). Similarly, its performance was sensitive to the resolution and the improved performance at a higher resolution was gained at the cost of overclustering of abundant cell types. In summary, aKNNO is able to identify both abundant and rare cell type accurately. As expected, aKNNO is superior than KNN in identifying rare cells and its high performance is less sensitive to the clustering resolution.

**Figure 2.**
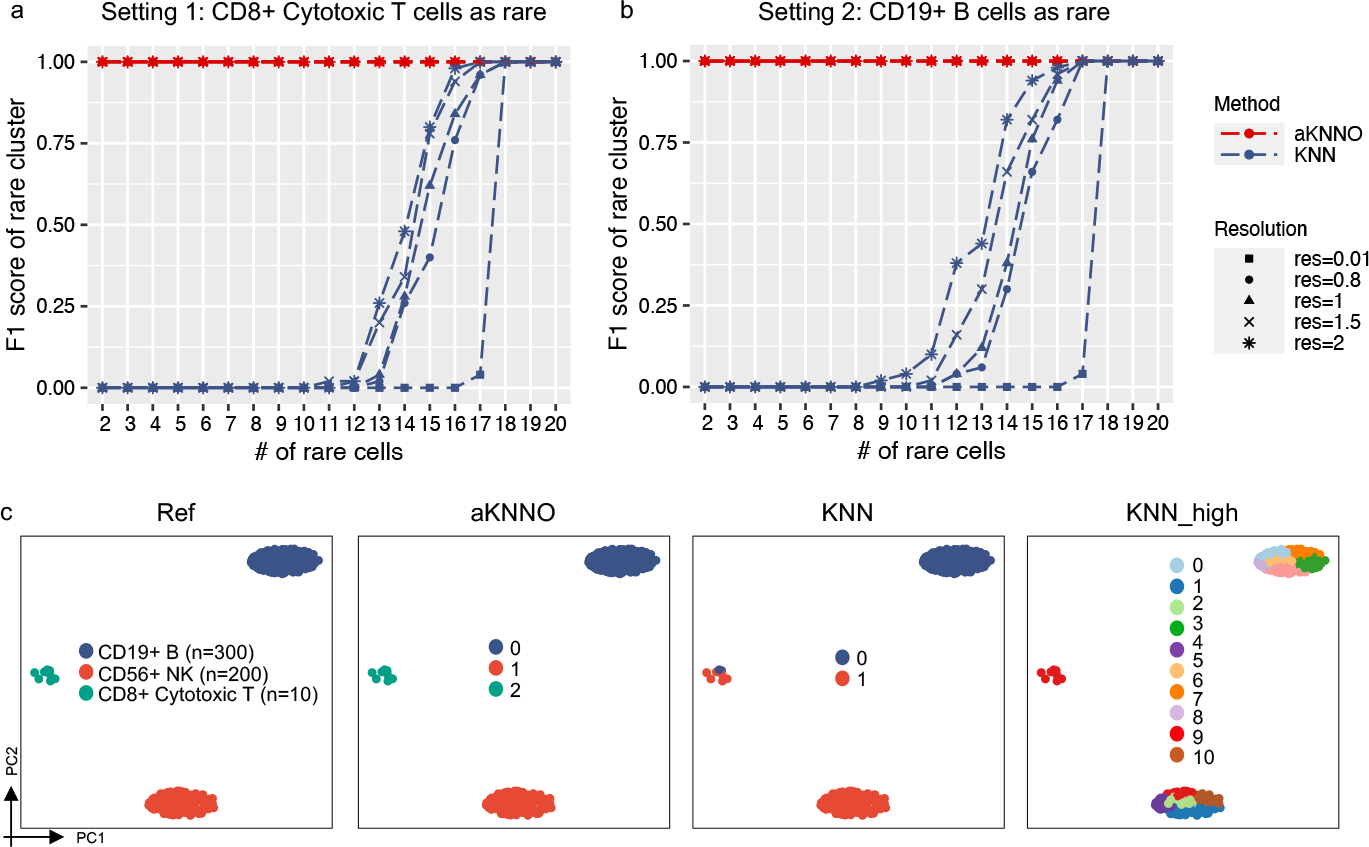
Application to simulated datasets generated from the PBMC68k dataset. F1 scores of aKNNO and KNN at different resolutions with the number of rare cells ranging from 2-20 in the first setting (a) and in the second setting (b). (c)The PCA plots labeled by the ground truth (Ref), labeled by aKNNO clustering at a resolution of 0.01 (aKNNO), labeled by KNN clustering at a resolution of 0.01 (KNN), and labeled by KNN clustering at a high resolution of 2 (KNN_high).

### Application to single-cell transcriptomics data from human pancreas

We applied aKNNO to a single-cell RNAseq dataset from human pancreas by inDrops technology, involving 5,542 cells and 14 cell populations manually annotated^23^. The 14 populations include three rare immune cell types, T, macrophage, and mast cells, and one rare epsilon cell type (Fig. 3a). aKNNO identified 25 clusters in total (Fig. 3b). aKNNO identified all the manually annotated cell types except schwann cells, including three rare immune cell types (clusters 18, 20, and 22 in Fig. 3b) and rare epsilon cells (cluster 23 in Fig. 3b, n=13). It also returned refined subclusters for abundant cell types. Compared to one ductal cell type in the manual annotation, aKNNO found three ductal clusters, cluster 5 (n=360), 15 (n=78), and 16 (n=58). The three clusters all expressed known ductal markers *KRT19* and *SOX9*, verifying their ductal identity. Compared to the abundant cluster 5, the cluster 15 was specifically positive for *OLFM4*, and the cluster 16 was high in *TFF1* and *IGFBP3* (Fig. 3e). These two clusters have been identified by a previous study on multipotent progenitor-like ductal cells, where *OLFM4*+ ductal was named as “transition to acinar 1” and *TFF1*+ *IGFBP3*+ was labeled as “activated/migrating progenitor cells”^24^. Besides, aKNNO identified two endothelial clusters with high expression of *PECAM1* and *PLVAP* (clusters 9 and 24). Compared to the cluster 9, the rare cluster 24 (n=10) showed highly specific expression of *PDPN* and *LYVE1* (Fig. 3e), which are well-known markers for lymphatic endothelial cells (LEC)^25^. As another example, aKNNO identified two activated stellate clusters with high level of *PDGFRB* (clusters 13 and 17), the abundant cluster 13 (n=106) with specific expression of *CXCL8* and *FGF2*, and the minor cluster 17 (n=38) with specific expression of *COL1A1* and *COL1A2* (Fig. 3e). This division of activated stellate cells have been reported in the original literature by further analyzing stellate cells^23^. In addition to discovering true rare cell clusters, aKNNO also detected rare doublets, clusters 19 (n=20) and 21 (n=17). They had relatively higher number of genes and UMIs than other corresponding cell types and showed signatures from two different cell types, indicating they are doublets. The cluster 19 not only showed high expression of beta cell markers like *INS*, but also highly expressed acinar cell markers like *CPA1*. The cluster 21 had both high expression of endothelial (*PECAM1* and *PLVAP*) and stellate markers (*PDGFRB*) (Supplementary Fig. S3).

**Figure 3.**
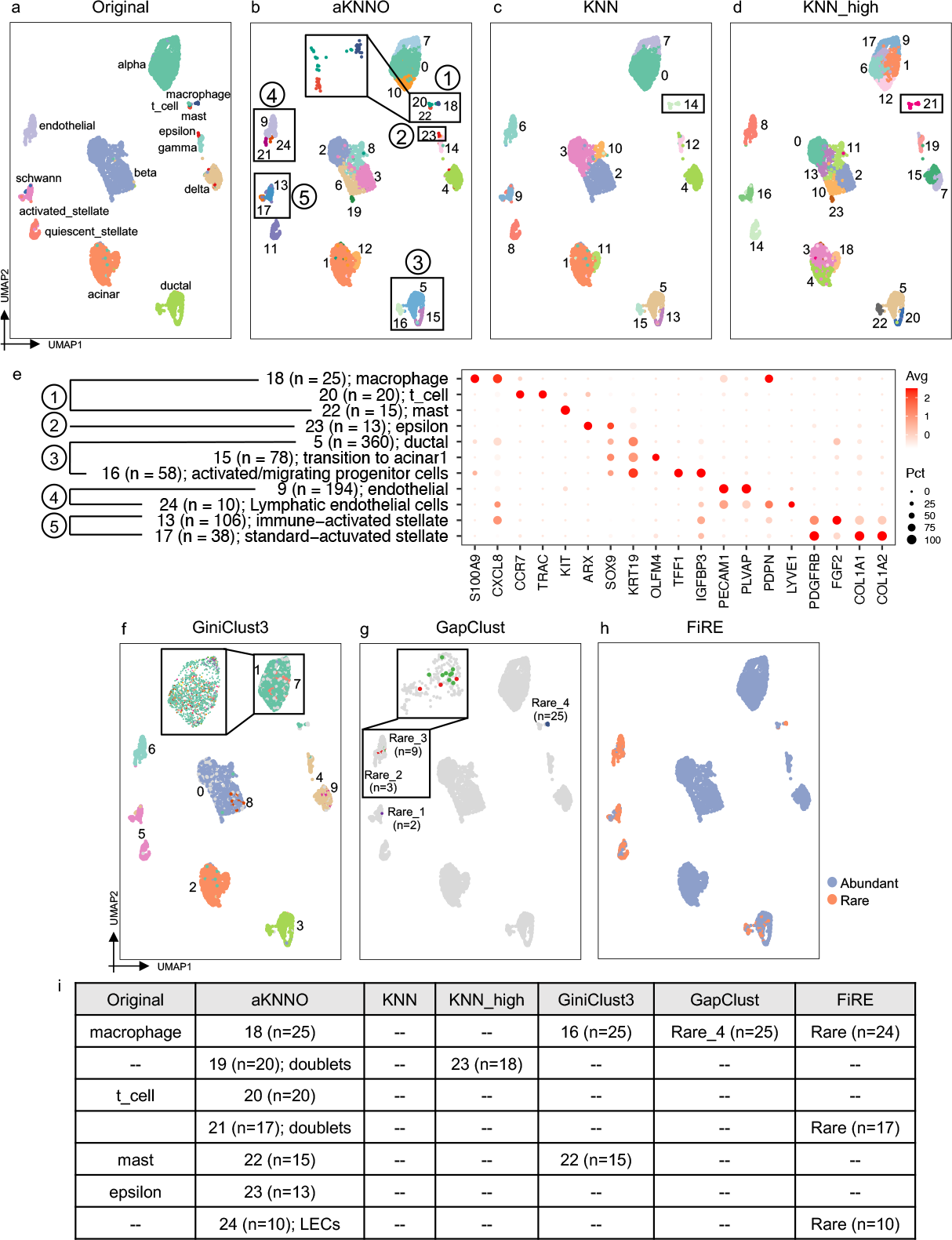
Application to single-cell RNAseq data from human pancreas. The UMAP plot labeled by the manual annotation from the original study (a), labeled by aKNNO clustering (b), KNN (c), KNN_high (d). (e) Dotplot of marker genes in clusters detected by aKNNO. (f) The UMAP plot labeled by the GiniClust3 result. Only the largest 10 clusters were shown since there were too many clusters. (g) The UMAP plot labeled by the GapClust result. (h) The UMAP plot labeled by the FiRE result. (i) A summary of rare clusters identified in the original study, aKNNO, KNN, KNN_high, GiniClust3, GapClust, and FiRE.

We compared aKNNO to KNN with default (r=0.8) and high resolutions (r=2, denoted as KNN_high). KNN discovered 16 clusters (Fig. 3c), while KNN_high identified 24 clusters in total (Fig. 3d). Surprisingly, KNN and KNN_high both failed to distinguish three rare immune clusters, T, macrophages, and mast cells (cluster 14 in the Fig. 3c and cluster 21 in the Fig. 3d), although they had very distinct features with high expression of *CCR7* and *TRAC* in T cells, *S100A9* and *CXCL8* in macrophages, and *KIT* in mast cells (Fig. 3e). KNN and KNN_high also missed the epsilon cells. KNN and KNN_high detected the two minor types of ductal cells, but were unable to identify lymphatic endothelial cells, the two activated stellate cells, and doublets (Figs. 3c and 3d). Although KNN_high obtained 24 clusters, it overclustered abundant cell types rather than found rare clusters. With 25 clusters, in contrast, aKNNO found many true rare cell types and doublets without overclustering those abundant ones. There was high agreement between aKNNO and KNN in clustering abundant cells (ARI>0.9), demonstrating that aKNNO is powerful in rare cell identification without scarifying its performance in abundant cells clustering.

We further compared aKNNO to three latest methods specifically designed or tailored to identify rare cell types, GiniClust3^18^, FiRE^19^, and GapClust^20^. GiniClust3 is an extension of GiniClust to identify both abundant and rare cell types, GapClust detects rare clusters, and FiRE quantifies rareness of each cell without clustering. aKNNO identified seven rare clusters with less than or equal to 25 cells (clusters 18-24, n=10∽25, Fig. 3i), where cells in each cluster were densely located together in the UMAP embedding. They were either manually annotated in the original literature (T, mast, macrophage, and epsilon cells), or supported by well-known marker genes (LEC, and two types of doublets) (Fig. 3i), demonstrating they were true rare cells or clusters. In comparison, GiniClust3 identified 85 clusters in total (Fig. 3f). Although GiniClust3 obtained so many clusters, it misclassified even activated and quiescent stellate cells into one group (cluster 5 in the Fig. 3f) and three types of ductal cells into one cluster (cluster 2 in the Fig. 3f), indicating its poor performance in clustering abundant cell types. For the rare cell clusters, GiniClust3 identified mast, and macrophage, but missed epsilon cells, lymphatic endothelial cells, and two types of doublets (Figs. 3f and 3i). Most rare clusters identified by GiniClust3 scattered and mixed well with other abundant cells in the UMAP embedding and not detected by other methods, suggesting they were highly likely to be false rare cells (Fig.3f). GapClust discovered four rare clusters, one of which is macrophage (n=25, Rare_4 in the Fig. 3g). The other three rare clusters containing two, three, and nine cells, respectively, were highly likely to be false positives since they scattered and mixed with endothelial and activated stellate cells in the UMAP embedding (Fig.3g). FiRE quantified 547 cells as being rare (Fig. 3h), which detected macrophage, lymphatic endothelial cells and one type of doublets correctly, but misidentified common endothelial cells and most of stellate cells as being rare (n>150), and most of T cells and mast cells (n<20) and all of epsilon cells (n=13) as being common (Figs. 3h and 3i). In summary, aKNNO identified more true and less false rare cells than GiniClust3, GapClust, and FiRE.

### Application to single-cell transcriptomics from mouse brain

We applied aKNNO to a single-cell RNA-seq dataset from mouse brain by 10x technology, involving 3,985 cells and 12 cell types manually annotated^26^. However, the UMAP embedding showed far more than 12 clearly-separated clusters, suggesting the original annotation is rough and imprecise (Fig. 4a). For example, there are two separated groups annotated as brain fibroblasts and multiple distinct clusters all labeled as microglia (Fig. 4a). aKNNO detected 29 clusters in total, which perfectly identified those separated groups in the UMAP as different clusters (Fig. 4b). For the two distinct groups manually annotated as brain fibroblasts, aKNNO identified one as the cluster 16 (n=41), and the other as the cluster 24 (n=17) (Fig. 4b). Both clusters expressed known fibroblast markers *Col1a1, Col1a2* and *Nupr1*, and the cluster 24 also specifically expressed *Fn1* and *Nov*, which was known as Fn1 fibroblasts^27^ (Fig. 4e). In comparison, KNN and KNN_high failed to distinguish these two types of fibroblasts (cluster 13 in the Fig. 4c and cluster 16 in the Fig. 4d). For the three distinct groups manually annotated as microglia, aKNNO detected six clusters (clusters 0,8,11, 17, 27, and 28 in the Fig. 4b). Five clusters had high expression of *Aif1*, a known marker of microglia, while the rare cluster 27 (n=7) had low expression of *Aif1* but specifically expressed *Xcl1, Cd3d* and *Cd3e* (Fig. 4e), suggesting its T cell identity. Each of the five types of microglia had specific gene expression signatures, cluster 0 (n=822) with high *Cx3cr1*, cluster 8 (n=117) with high *Ccl4*, cluster 11 with high *Apoe* (n=61), rare cluster 17 (n=36) with high *Cd74*, and rare cluster 28 (n=5) with high *Ifit3* and *Isg15* (Fig. 4e). The microglia clusters 0, 8, 11, and 17 have different functions reported by previous studies^28,29^, and the cluster 28 is highly enriched in IFN-response genes, suggesting it is a novel type of microglia. Among the five types of microglia and T cells, KNN only identified three and KNN_high found five types (Figs. 4c and Figs. 4d). Besides GABAergic and Glutamatergic neurons, aKNNO identified two more types of rare neurons (clusters 21 and 23 in the Fig. 4b, n=23 and n=18 respectively), where the cluster 21 had specific expression of *Tle4* and *Rprm* and the cluster 23 was positive for *Nxph1* and *Nxph3* (Fig. 4e). Those genes all play important roles in neurons^30^, suggesting they are novel neuron types. KNN failed to identify both types of neurons, while KNN_high missed one type. In addition, AKNNO discovered two rare and distinct clusters (the cluster 20, n=26; the cluster 26, n=10 in the Fig. 4b), which were annotated as “multiplets” in the original annotation. The cluster 20 had high expression of *Alas2* and *Hbb-bt*, which are erythroid markers. The cluster 26 had specific expression of *Otx2*, which is known to regulate progenitor identity and neurogenesis in the midbrain^31,32^(Fig. 4e). Besides, aKNNO and KNN were in high agreement on their clustering of abundant cells (ARI>0.92), demonstrating aKNNO doesn’t reduce their power at clustering abundant cells when it achieves great performance in identifying rare ones.

**Figure 4.**
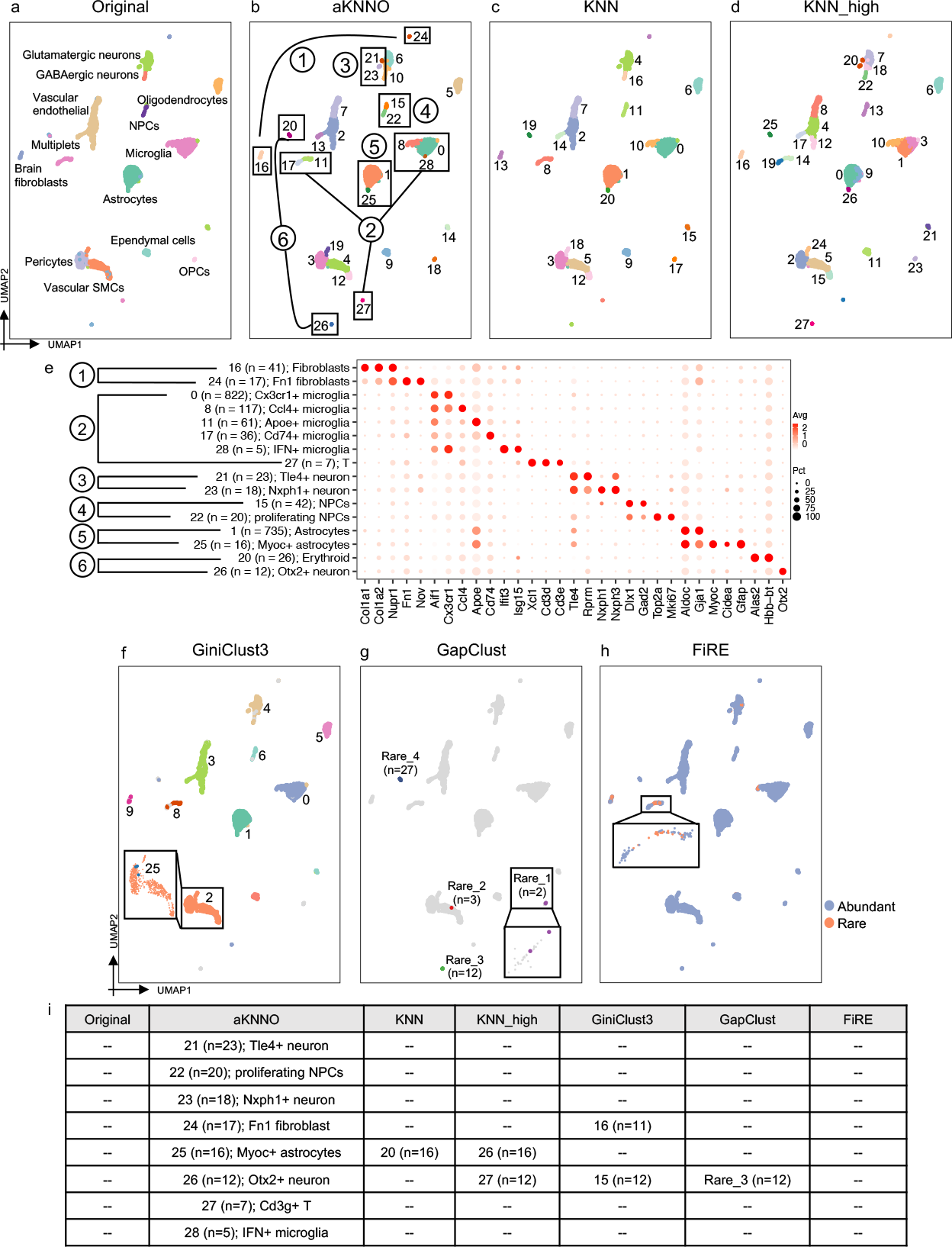
Application to single-cell RNAseq data from mouse brain. The UMAP plot labeled by the manual annotation from the original study (a), aKNNO (b), KNN (c), KNN_high (d). (e) The Dotplot of marker genes in clusters detected by aKNNO. (f) The UMAP plot labeled by the GiniClust3 result. Only the largest 10 clusters were shown. (g) The UMAP plot labeled by the GapClust result. (h) The UMAP plot labeled by the FiRE result. (i) A summary of rare clusters identified in the original study, aKNNO, KNN, KNN_high, GiniClust3, GapClust, and FiRE.

We further compared rare cells identified by aKNNO, GiniClust3, FiRE, and GapClust. aKNNO identified eight rare clusters with less than 25 cells (clusters 21-28, n=5∽23) (Figs. 4b and 4i). As discussed above, cluster 21 was *Tle4*+ neuron, cluster 23 was *Nxph1*+ neuron, cluster 24 was *Fn1* fibroblast, cluster 26 was Otx2+ neuron, cluster 27 was T cell, and cluster 28 was IFN+ microglia. Cluster 22 was similar to cluster 10 with high expression of *Dlx1* and *Gad2*, which was manually annotated as neural progenitor cell (NPCs). However, cluster 22 expressed high *Top2a* and *Mki67* (Fig. 4e), suggesting it was proliferating NPCs. Cluster 25 and cluster 1 both had high expression of known astrocytes markers, such as *Aldoc* and *Gja1*. Cluster 25 had specific expression of *Myoc, Cidea* and *Gfap* (Fig. 4e), which has been reported as a new type of Astrocytes^33^. In comparison, GiniCluster3 identified 27 clusters in total (Fig. 4f). For abundant cell types, it misidentified pericytes and vascular SMCs as one group (cluster 2 in the Fig. 4f), indicating its inefficiency in large cluster detection. Among the eight rare cell types identified aKNNO, GiniClust3 only identified Fn1 fibroblast and Otx2+ neuron (Fig. 4i). Cells in other rare clusters by GiniClust3 scattered in the UMAP embedding and mixed with other abundant cells, indicating they are not true rare cells (Fig. 4f). GapClust obtained four rare clusters, containing 2, 3, 12, and 27 cells, respectively (Fig. 4g). The cluster with 12 cells (Rare_3 in the Fig. 4g) corresponded to the cluster 26 of aKNNO, which is Otx2+ neuron. The cluster with 27 cells (Rare_4 in the Fig. 4g) matched the cluster 20 in the aKNNO, which was not that rare compared to the eight rare clusters. The other two rare clusters mixed with abundant cells in the UMAP embedding (Fig. 4g), suggesting they are not real rare. FiRE only found 56 rare cells (Fig. 4h), most of which mixed with microglia and fibroblasts. It failed to detect any of the eight rare clusters in aKNNO (Fig. 4i). Consistent with the results from human pancreas, aKNNO achieved higher sensitivity and specificity than other methods in identifying rare cells.

### Application to spatial transcriptomics data from mouse posterior brain

Spatial transcriptomics map out organizational structures of cells along with their transcriptomics profiles, providing powerful tools for understanding spatial and functional arrangement of tissues^34^. We applied aKNNO to a 10x Visium dataset generated from mouse sagittal posterior brain, which has complicated tissue structures. We used the brain anatomical reference annotations from the Allen Mouse Brain Atlas and H&E image as the ground truth (Figs. 5a and 5e). aKNNO identified 31 clusters from the 3,355 spots, which stereotyped anatomical structures precisely (Fig. 5b). For example, the hippocampal system consists of the dentate gyrus (DG), cornu ammonis (CA) fields that are subdivided into four regions (CA1-CA4), and the subiculum^35^. The brain anatomical reference annotations and the H&E image of the data show dorsal and ventral hippocampus, where the dorsal contains DG, CA3 and CA1, and the ventral includes DG, CA3 and subiculum (highlighted in red boxes in the Figs. 5a and 5e). Previous transcriptomics studies on neuronal classes of the hippocampus found not only cell-class-but also region-specific-expression profiles, indicating heterogeneous populations along the dorsal-ventral axis^36^. aKNNO discovered six clusters successfully with known cell- and region-specific expression (clusters 21, 22, 25, 26, 27, and 30 in the Fig. 5g), which mapped to the six anatomical patterns precisely. In the ventral hippocampus, cluster 30 (n=13) aligned exclusively to DG and had high expression of known DG-specific (*Prox1*) and ventral-specific genes (*Trhr*)^36^ (Fig. 5F). Cluster 27 (n=17) depicted CA3 with CA3-specific expression of *Ociad2* and *Cpne7*^36^, and cluster 22 corresponded to subiculum with known cell-class and region-specific expression of *Dio3*^37^ (Fig. 5f). In the dorsal region, cluster 25 (n=31) lined up with DG with DG-specific gene *Prox1* and dorsal-specific gene *Lct*^36^, Cluster 26 (n=20) mapped to CA3 with high CA3_dorsal-specific expression of *Iyd* ^36^, and cluster 21 (n=54) matched CA1 with high expression of *Fibcd1*^36^ (Fig. 5f). In comparison, KNN and KNN_high discovered 15 and 28 clusters, respectively, which showed poorer performance in defining anatomical patterns than aKNNO (Figs. 5c and 5d). For example, KNN and KNN_high failed to distinguish the six hippocampus subpopulations. KNN only found two clusters, where it misclassified DG_ventral and the whole dorsal hippocampus into one cluster, and it miscategorized CA3_ventral and subiculum into one and also masked them into the surrounding region (Fig. 5h). KNN_high detected four clusters, where it misclassified DG_ventral, CA3_dorsal, and DG_dorsal into one cluster, and it failed to distinct CA1_dorsal, CA3_ventral and subiculum from their surrounding regions (Fig. 5i).

**Figure 5.**
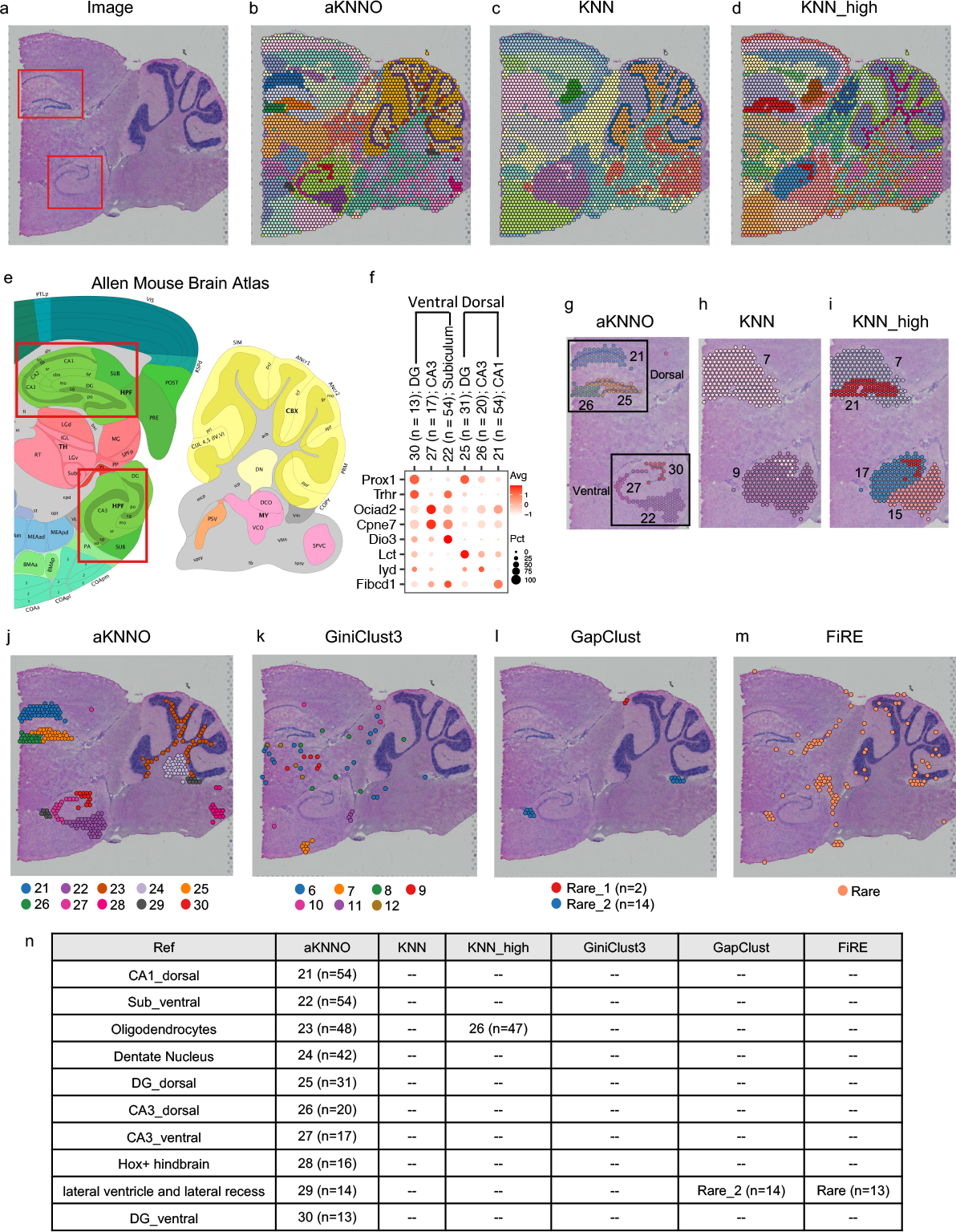
Application to 10x Visium spatial transcriptomics data from mouse posterior brain. (a) H&E image. The spatial plot annotated by aKNNO (b), KNN (c) and KNN_high (d). (e) Allen Brain Institute reference atlas diagram. (f) Dotplot of marker genes in the dorsal and ventral hippocampus structures. The spatial plot focusing on dorsal and ventral hippocampus structures annotated by the six cell clusters identified by aKNNO (g), two cell clusters detected by KNN (h), three cell clusters by KNN_high (i). The spatial plot of rare cell clusters detected by aKNNO (j), GiniClust3 (k), GapClust (l), and FiRE (m). (n) A summary of rare clusters identified by aKNNO, KNN, KNN_high, GiniClust3, GapClust, and FiRE.

The ten rare clusters identified by aKNNO (clusters 21-30 in the Fig. 5j, n<55) all aligned exclusively to a specific brain region with functional meaning, including CA1_dorsal (cluster 21, n=54), subiculum_ventral (cluster 22, n=54), Oligodendrocytes in the white matter (cluster 23, n=48), Dentate Nucleus (cluster 24, n=42), DG_dorsal (cluster 25, n=31), CA3_dorsal (cluster 26, n=20), CA3_ventral (cluster 27, n=17), Hox genes-enriched hindbrain (cluster 28, n=16), lateral ventricle and lateral recess (cluster 29, n=14) and DG_ventral (cluster 30, n=13) (Figs. 5e, 5j and 5n). In comparison, GiniClust3 detected only 13 clusters, which agreed poorly with the Allen Brain Institute reference atlas diagram (Supplementary Fig. S4). For example, multiple cortical layers were misclassified into one cluster and none of the hippocampus structure were identified correctly (Supplementary Fig. S4). The rare clusters (clusters 6-12 in the Fig. 5k with n<55) scattered randomly in the spatial region and did not map to specific anatomical regions, suggesting that they were highly likely to be false. None of the ten rare clusters in aKNNO were detected by GiniCluster3 (Fig. 5n). GapClust only identified two rare clusters (Fig. 5l). One cluster containing 14 cells aligned to lateral ventricle and lateral recess region (cluster 29 in the aKNNO), while the other cluster with two cells was identified to be common by other methods. FiRE identified 105 rare cells (Fig. 5m). Among them, 13 cells were lined up with lateral ventricle and lateral recess region (cluster 29 in the aKNNO). 66 cells were smooth muscle cells with high level of *Ogn* and *Prdm6*, which were also identified by aKNNO (Supplementary Fig. S5, n=105) and were not that rare compared to the ten rare clusters. Other cells scattered in the brain region and might not be true rare cells. In summary, the ten rare clusters identified by aKNNO stereotyped subtle anatomical structures, whereas GapClust and FiRE only recovered the lateral ventricle and lateral recess structure (Fig. 5n).

Recently, several integrative approaches combining expression, spatial location and histology images have been developed to improve the performance of spatial transcriptomics clustering^38-41^. We compared aKNNO with four integrative approaches, stLearn^39^, SpaGCN^38^, GraphST^40^, and BayesSpace^41^. We set their number of clusters to 31, matching the cluster number of aKNNO (Figs. 6a-6e). Using the fine-grained hippocampus structures as an example, aKNNO stereotyped six anatomical patterns precisely (Fig. 6f). In contrast, all the four integrative approaches failed to resolve the six patterns. BayesSpace misclassified CA3_dorsal and CA1_dorsal into one (cluster 28 in the Fig. 6g), DG_dorsal and DG_ventral into one (cluster 26 in the Fig. 6g), and also failed to distinguish CA3_ventral and subiculum from their surrounding regions (clusters 3 and 23 in the Fig. 6g). GraphST identified CA1_dorsal (cluster 22 in the Fig. 6h), but misrecognized DG_dorsal and DG_ventral into one (cluster 24 in the Fig. 6h), and CA3_dorsal and CA3_ventral into one (Cluster 10 in the Fig. 6h). SpaGCN successfully identified CA3_dorsal (cluster 27 in the Fig. 6i) and CA3_ventral (cluster 23 in the Fig. 6i), but misclassified DG_dorsal and DG_ventral into one (cluster 30 in the Fig. 6i) and also lost the subiculum structure. stLearn, similar to GraphST, detected CA1_dorsal (cluster 21 in the Fig. 6j), but failed to distinguish between CA3_dorsal and CA3_ventral (cluster 18 in the Fig. 6j) and between DG_dorsal and DG_ventral (cluster 27 in the Fig. 6j). The comparison demonstrated that aKNNO, using gene expression alone, resolved tissue structures more accurately than those integrative approaches.

**Figure 6.**
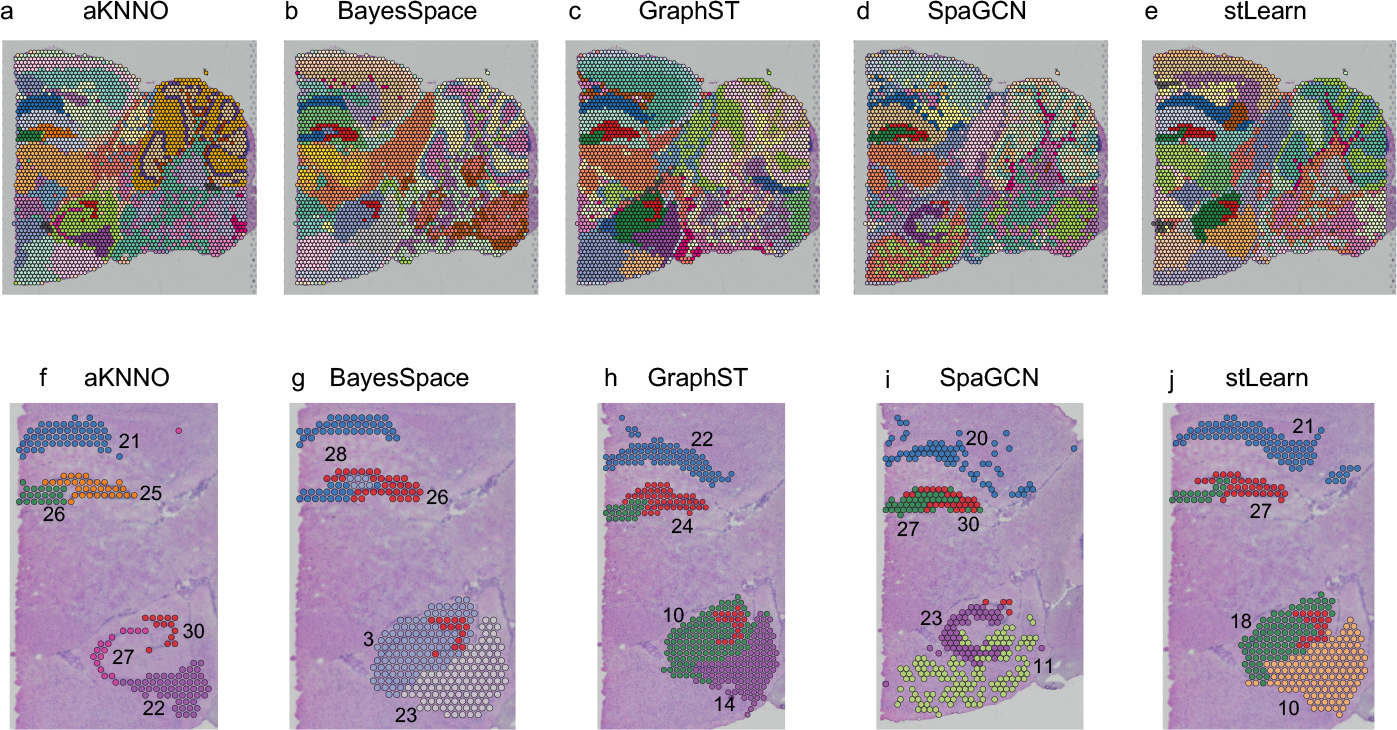
Comparison between aKNNO, BayesSpace, GraphST, SpaGCN, and stLearn on the 10x Visium spatial transcriptomics data from mouse posterior brain. The spatial pot annotated by aKNNO (a), BayesSpace (b), GraphST (c), SpaGCN(d), and stLearn(e). Detailed view of clustering in hippocampus structures in aKNNO (f), BayesSpace (g), GraphST (h), SpaGCN(i), and stLearn (j).

## Discussion

The accurate detection of abundant and rare clusters simultaneously is crucial for characterizing cellular heterogeneity in single-cell and spatial transcriptomics analysis. Here, we presented aKNNO, an adaptive *k*-nearest neighbor graph with optimization for the community-detection-based clustering. Compared to traditional *k*NN specifying an universal *k* for all cells, aKNNO chooses *k* adaptively for each cell based on its local distance distribution. The adaptive strategy assigns a small *k* for rare cells and a large *k* for abundant cells, enabling to capture the inherent cellular structure accurately. aKNNO has been extensively evaluated on 38 simulated scenarios and 20 single-cell and spatial transcriptomics data from different species, tissues and technologies. Additional analysis on mouse intestine from 10x, organoids from CEL-seq, mouse habenula from inDrops, spatial main olfactory bulb, spatial coronal posterior brain, and 12 pan-cancer datasets were included in the Supplementary Material. The results consistently demonstrated that aKNNO outperformed traditional *k*NN-based approach in identifying abundant and rare cell types in a single run. aKNNO was also far more superior than those methods specifically designed or tailored for rare cells detection in terms of both sensitivity and specificity.

aKNNO provides an optimization step to tune *σ*, a hyperparameter controlling the sensitivity to local distance change. High sensitivity to local distance change would result in many localized subclusters and lead to overclustering, while low sensitivity would lose the adaptive ability and reduce to the traditional *k*NN. aKNNO uses grid search to find the optimal *σ* for each dataset. We compared the performance between the optimal and the default *σ* (−0.5). We found the optimal *σ* was different across datasets. aKNNO using the optimal *σ* was able to find more true rare cell types than the approach using the default *σ* without optimization (Supplementary Figs. S6-S9). That is, the optimization step did help balance between sensitivity and specificity in rare cells identification. aKNNO is less sensitive to the clustering resolution than *k*NN-based approaches thanks to its ability to capture the inherent cellular structure more accurately. For the *k*NN-based approaches, the choice of the resolution has a large effect on the outcome and there is no consensus how to find the right resolution. In our evaluation, some rare clusters missed at a low resolution (KNN) might be detected at a high resolution (KNN_high). In contrast, rare clusters were identified by aKNNO at the default resolution and remained stable when the resolution increased.

aKNNO builds a nearest neighbor graph, which can be seamlessly incorporated into existing analysis pipelines (see tutorials in the GitHub) and also used for any graph-based clustering approaches, such as Louvain clustering, Spectral clustering, Leiden clustering, and Minimal Spanning Tree. aKNNO not only identified true rare clusters, but also found doublets and empty droplets. For example, aKNNO found two types of doublets in the human pancreas and one cluster with only two empty droplets in the mouse habenula dataset. That is, aKNNO can be used to remove poor quality cells that are not eliminated completely in the preprocessing step. Although an adaptive strategy is included to build the NN-graph, the computational burden added is neglectable compared to the traditional *k*NN-based approach. The runtime of aKNNO and KNN is almost the same especially when there are large number of cells (Supplementary Fig. S10).

Spatial transcriptomics is an emerging technology that provides a roadmap of transcriptional activity within tissue sections. To better decipher domains or cell types that are spatially coherent in both gene expression and histology, a number of integrative approaches to combine gene expression, spatial location, histology, and H&E image have been developed^38-41^. Integrating multi-modal information are expected to define cell types or domains accurately than using gene expression alone. Surprisingly, aKNNO is able to stereotype anatomical cell types precisely using gene expression alone. For example, the six clusters identified by aKNNO aligned almost perfectly with the six dorsal and ventral hippocampus structures (DG_dorsal, DG_ventral, CA3_dorsal, CA3_ventral, CA1_dorsal, and subiculum_ventral). Each common and rare cluster detected by aKNNO mapped to specific regions in the Allen Brain Institute reference atlas diagram, suggesting they are functionally meaningful. Some clusters or spatial regions were even not identified by those integrative approaches (Fig. 6 and Supplementary Material). The results suggest that gene expression alone is enough to define spatial cell types or domains when the appropriate strategy is used.

## Methods

### Single-cell and spatial transcriptomics data processing

Single cell RNAseq datasets were filtered and processed by the Seurat package^15^. Specifically, cells expressing less than 200 genes and genes expressed in less than three cells were excluded. Data were normalized to the 10,000 UMI and the top 2,000 highly variable genes were selected by the vst method. Principal component analysis (PCA) was used to reduce dimension and UMAP embedding was computed for visualization.

Spatial transcriptomics data were processed by the Seurat package^42^. Particularly, data were normalized and the top 3,000 highly variable genes were selected using SCTtransform^43^. Genes expressed in less than a certain percentage of all spots were excluded, which was set to 10%, 0%, and 0% for the mouse coronal and sagittal posterior brain, and main olfactory bulb, respectively. Principal component analysis (PCA) was used to reduce dimension and UMAP embedding was computed for visualization.

### aKNNO

aKNNO includes five steps: 1) calculate the distances within the Kmax nearest neighbors for all cells; 2) choose the actual number of the true nearest neighbors *k* adaptively for each cell to construct an adaptive *k*NN graph; 3) build the shared nearest neighbor graph; 4) Louvain clustering on the shared neighbor graph; 5) repeat steps 2)-4) for optimization. (Fig. 1).

1. calculate the distances within the Kmax nearest neighbors for all cells; The top 50 PCs were used to calculate the distances between cells. Given a maximum number of the nearest neighbors Kmax, RANN package is used to find the Kmax nearest neighbors of all cells and calculate the distances.
2. construct an adaptive *k*NN graph by choosing *k* adaptively for each cell; For each cell *i*, the distance to its Kmax nearest neighbors are sorted in an ascending order (d*i*1<d*i*2<…d*i*Kmax) and *k* is chosen based on the Kmax distance distribution (Cai et al. 2022)^44^. Several algorithms have been proposed to find adaptive neighbors^45-48^. We used the algorithm described in Cai et al. (Cai et al. 2022)^44^, which does not require the regularization term and thus be more computational efficient. Assuming there are *n* cells in total (1,2,…n), for the cell *i*, the cell *j* can be connected to *i* as a neighbor with probability S*ij*. Intuitively, a smaller distance should be assigned a larger probability. Cai et al. (Cai et al. 2022) computes the probability by minimizing the following objective function:

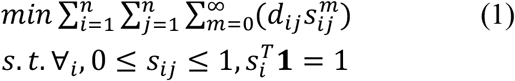

where m is the number of power, *d*_*ij*_ is the distance between the cell *i* and *j*. The standard Karush-Kuhn-Tucker conditions are used to solve the equation (1). The optimal solution is:

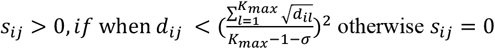

where *σ* is a hyperparameter to control the sensitivity to the local distance change (*σ* ≤ 0). *s*_*ij*_ > 0 indicates there is a connection between *i* and *j*, while *s*_*ij*_ = 0 means no connection. Therefore, for the Kmax nearest neighbors of the cell *i*(d_*i*1_<d_*i*2_<d_*i*Kmax_), if 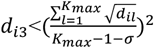 but 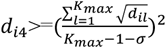, then the true nearest neighbor *k* for the cell *i* is set to 3 automatically (*k*=3). In this way, aKNNO chooses the *k* adaptively based on the local distance distribution for each cell.
3. build the shared nearest neighbor graph; The adaptive *k*NN graph is reweighted based on the shared nearest neighbors of each pair of cells, which make it more robust to outliers and noise. The Jaccard similarity is used to build the shared nearest neighbor graph.
4. Louvain clustering on the shared neighbor graph; The Louvain clustering in the Seurat package is used to group cells on the shared nearest neighbor graph.
5. repeat steps 2)-4) for optimization. *σ* is a hyperparameter to control the sensitivity to the local distance change (default=-0.5). A too negative *σ* would make aKNNO very sensitive to local distance change and lead to overclustering, while aKNNO would loose the adaptive ability if *σ* is not negative enough. To find the optimal *σ*, aKNNO employs a grid search to find the *σ* that balances between the sensitivity and specificity. aKNNO decreases *σ* in each repeat and choose the *σ* before there is a rapid increase in the number of communities detected, which suggests overclustering (Fig. 1).

### Comparison with other methods

We compared aKNNO with the conventional *k*NN-based clustering method in the Seurat package (denoted as KNN). aKNNO identified clusters at the default solution (r=0.8), while KNN found clusters at the default resolution (r=0.8) and a high resolution (r=2, denoted as KNN_high). To make a fair comparison, aKNNO and KNN processed the data exactly the same way, including the highly variable genes selection, the number of PCs to calculate the distance, the method to find the nearest neighbors, and Louvain clustering. The only difference between aKNNO and KNN is the graph used for clustering. aKNNO generates clustering from the adaptive *k*NN graph, while KNN from the traditional *k*NN graph.

We also compared aKNNO with three methods specifically designed or tailored to identify rare cell types, GiniClust3^18^, GapClust^20^, and FiRE^19^. GiniClust3 is an extension of GiniClust, which discovers rare cells based on genes with high Gini index. FiRE uses the Sketching technique to assign a rareness score to each cell^19^. GapClust captures the abrupt local distances change to find rare cell clusters^20^. GiniClust3 identifies abundant and rare clusters simultaneously, GapClust finds rare clusters only, and FiRE quantifies rareness of each cell without clustering. Default parameters were used to run GiniClust3, GapClust and FiRE. GapClust requires a normalized expression matrix as the input, therefore the data were normalized to 10,000 UMI without log-transformation for GapClust.

In the spatial transcriptomics data, we further compared aKNNO with four integrative approaches, BayesSpace^41^, GraphST^40^, SpaGCN^38^, and stLearn^39^. BayesSpace implements a full Bayesian model that uses the information from spatial neighborhoods for resolution enhancement for spatial clustering. stLearn first normalizes gene expression by distance measures on morphological similarity and neighborhood smoothing and then clustering cell types from the normalized expression profiles. SpaGCN uses a graph convolutional network that integrate gene expression, spatial location and histology images. GraphST is a graph self-supervised contrastive learning method that combines spatial location and expression. Default parameters were used to run BayesSpace, GraphST, SpaGCN, and stLearn. The number of clusters was set to the same as aKNNO.

### Simulation studies

We generated simulated datasets by sampling three cell types from the PBMC 68k dataset^22^, CD19+ B, CD56+ NK and CD8+ Cytotoxic T. We created two settings. In one setting, the rare cell type was similar to one of the abundant cell type, i.e., rare CD8+ Cytotoxic cells but abundant CD19+ B (n=300) and CD56+ NK (n=200) cells. In the other setting, the rare cell type was distinct from both abundant cell types, i.e., rare CD19+ B cells but abundant CD56+ NK (n=200) and CD8+ Cytotoxic T (n=300). In each setting, we simulated 19 scenarios with the number of rare cells ranging from 2 to 20. In each scenario, 50 simulated datasets were generated by random sampling the three cell types from the PBMC 68k dataset.

## Code and Data Availability

The aKNNO R package is freely available in the GitHub repository https://github.com/liuqivandy/aKNNO). The single cell RNAseq data for human pancreas, mouse brain, mouse intestine, intestinal organoids, and mouse habenula are available at the GEO under accession codes GSE84133, GSM3580745, GSM4521364, GSE62270, GSM4411753, respectively. The 10x Visium data for mouse coronal and sagittal posterior brain, and main olfactory bulb are downloaded from the 10x Genomics website (https://www.10xgenomics.com/resources/datasets). The single cell RNAseq dataset of the melanoma is obtained from GEO under the accession id GSE72056 and the other 11 cancer datasets were downloaded from the database Tumor Immune Single-cell Hub 2 (TISCH2)^49^. The code to reproduce the results in the manuscript is also available in the GitHub as tutorials.

## Supplementary data

Supplementary data are available online.

## Author Contribution

Q.L. and Y.S. conceived and designed the study. J.L. and Q.L. collected and analyzed the sequencing data. J.L. and Q.L. implemented the algorithm and developed the package. J.L. and Q.L. wrote the manuscript. All authors read and approved the final manuscript.

