## Supplemental figures for "Single-cell and Spatial Transcriptomics Clustering with an Optimized Adaptive K-Nearest Neighbor Graph"

### Additional file 1

**Supplementary Fig. S1.** The highest adjusted rand index of aKNNO (red) and KNN (blue) with the ground truth in the first setting (a) and in the second setting (b). The adjusted rand index at each resolution was calculated and the highest value was selected.

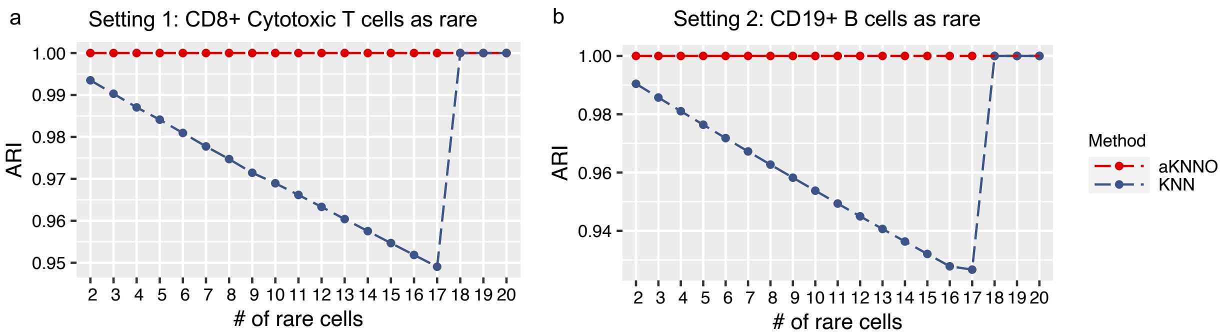

**Supplementary Fig. S2.** The performance of aKNNO, KNN, and KNN_high on a simulated dataset with 17 rare CD8+ cytotoxic T cells, 200 CD56+ NK, and 300 CD19+ B cells.

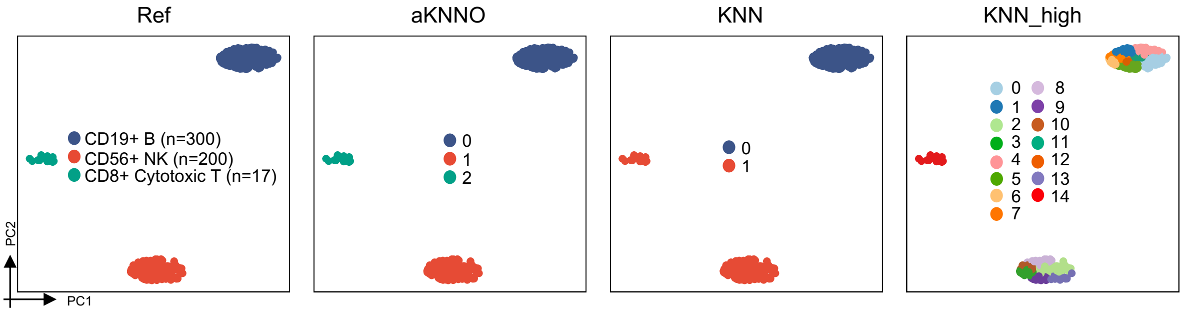

**Supplementary Fig. S3.** The distribution of UMI counts, the number of genes, gene signatures in the cluster 19 and the cluster 21 identified by aKNNO in the human pancreas data. The cluster 19 (a) and the cluster 21 (b) had both higher UMI counts and number of genes than other corresponding clusters. (c) The cluster 19 expressed high beta (*INS*) and acinar markers (*CPA1*). The cluster 21 had both high expression of endothelial (*PECAM1* and *PLVAP*) and stellate markers (*PDGFRB* and *RGS5*).

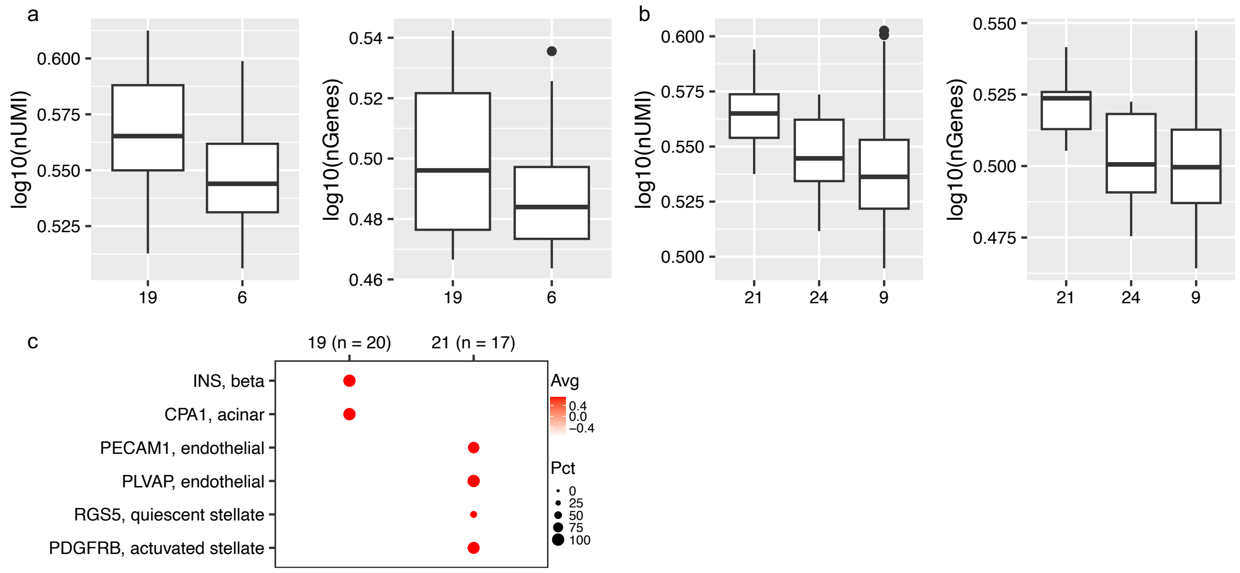

**Supplementary Fig. S4.** GiniCluster3 clustering on the 10x Visium mouse posterior brain

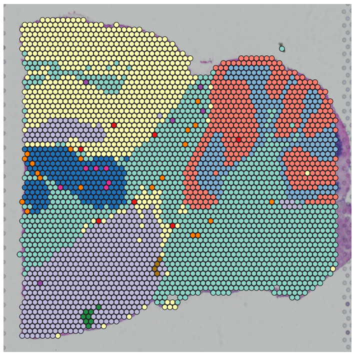

**Supplementary Fig. S5.** (a) The cluster 16 identified by aKNNO. (b) Spatial expression plots of *Ogn* and *Prdm6* in the 10x Visium mouse posterior brain data.

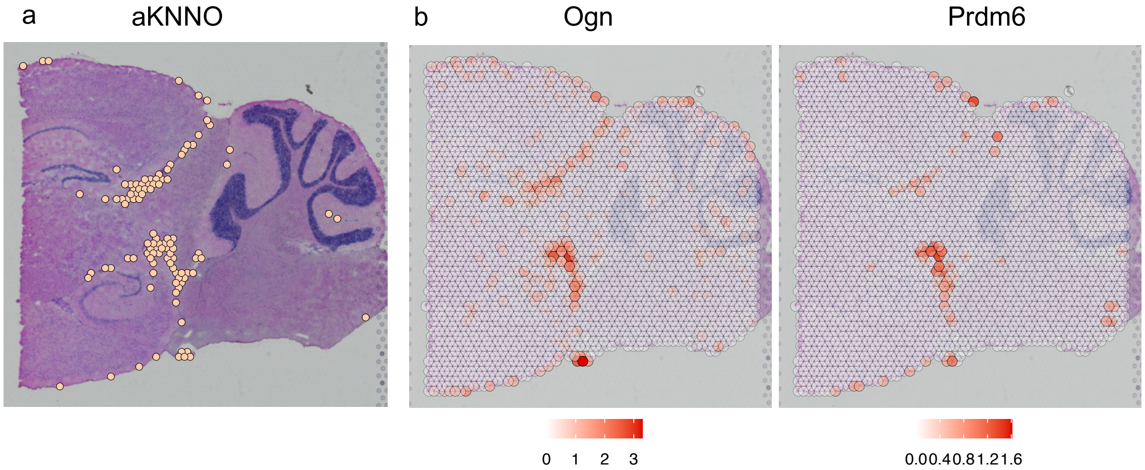

**Supplementary Fig. S6.** Performance of aKNNO and aKNNO without optimization at the default delta of -0.5 on the human pancreas data. (a) The number of communities and singletons detected at different delta values. The optimal delta of -0.8 (dash line) was chosen automatically. (b) Clustering of aKNNO. (c) Clustering of aKNNO at delta of -0.5 without optimization.

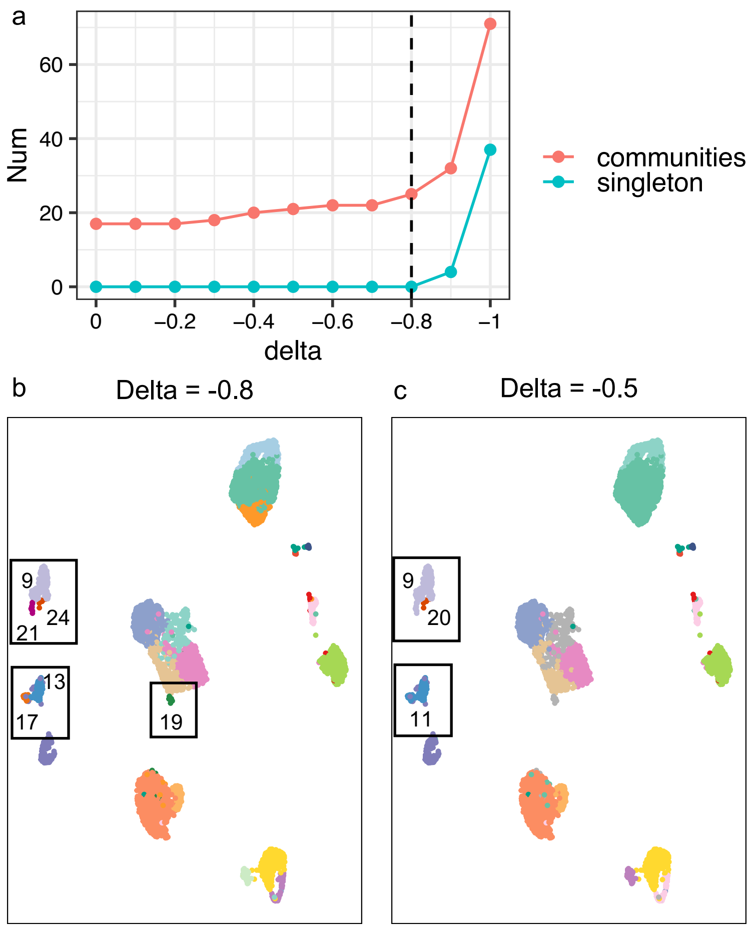

**Supplementary Fig. S7.** Performance of aKNNO and aKNNO without optimization at the default delta of -0.5 on the mouse brain data. (a) The number of communities and singletons detected at different delta values. The optimal delta of -0.8 was chosen automatically. (b) Clustering of aKNNO. (c) Clustering of aKNNO at delta of -0.5 without optimization.

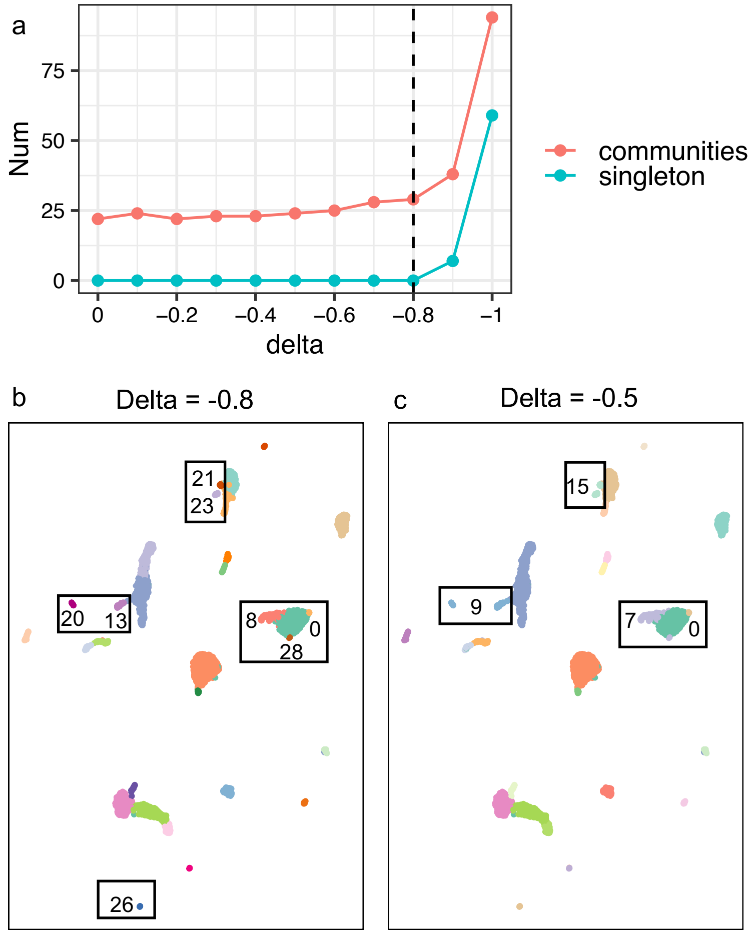

**Supplementary Fig. S8.** Performance of aKNNO and aKNNO without optimization at the default delta of -0.5 on the mouse intestine data. (a) The number of communities and singletons detected at different delta values. The optimal delta of -0.9 was chosen automatically. (b) Clustering of aKNNO. (c) Clustering of aKNNO at delta of -0.5 without optimization.

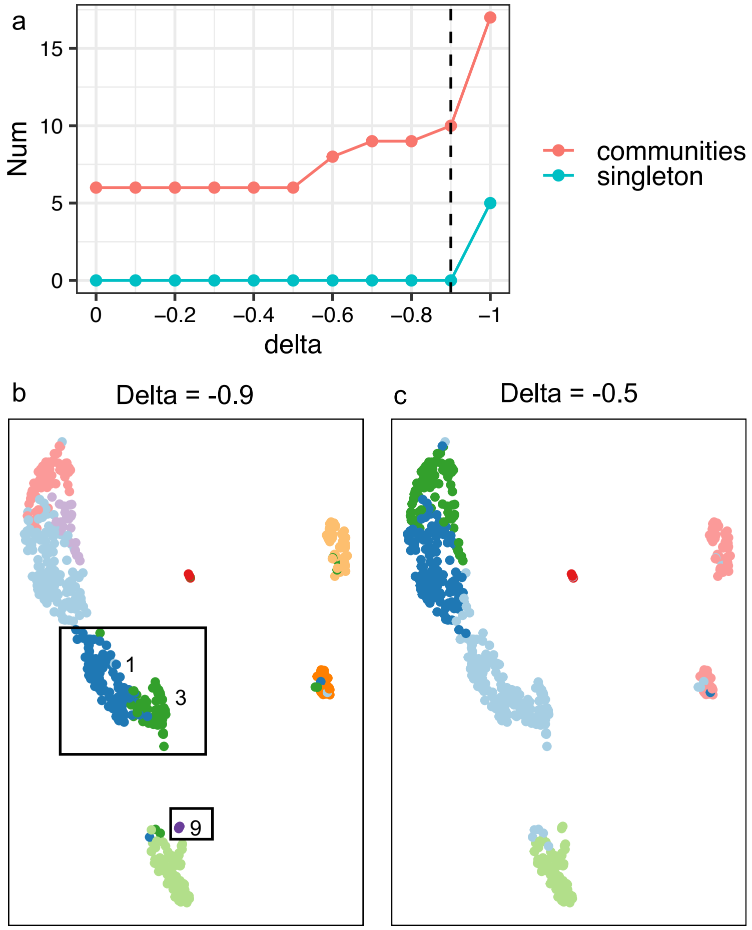

**Supplementary Fig. S9.** Performance of aKNNO and aKNNO without optimization at the default delta of -0.5 for the 10x Visium mouse posterior brain data. (a) The number of communities and singletons detected at different delta values. The optimal delta of -0.9 was chosen automatically. (b) Results of aKNNO. (c) Results of aKNNO at delta of -0.5 without optimization.

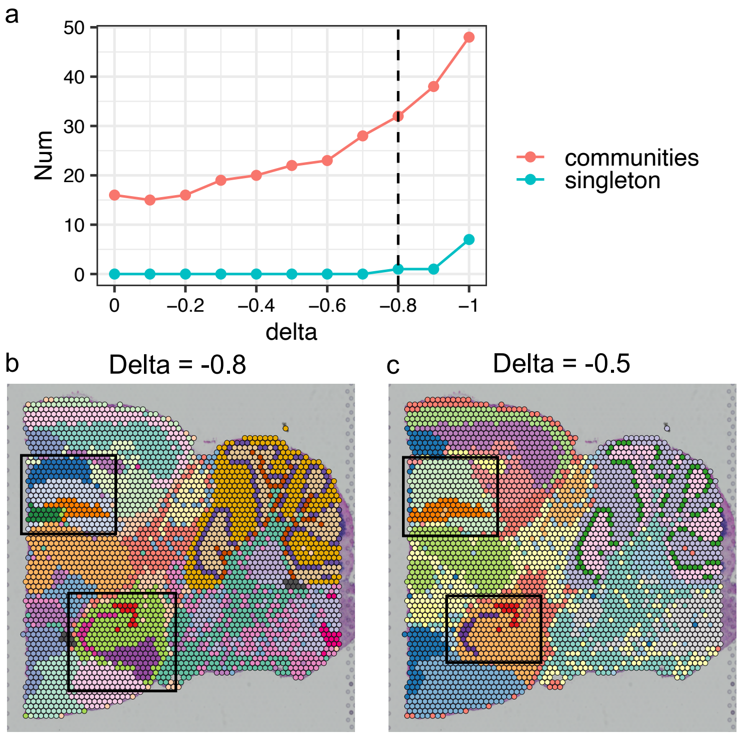

**Supplementary Fig. S10.** Runtime of aKNNO and KNN as the number of cells increases.

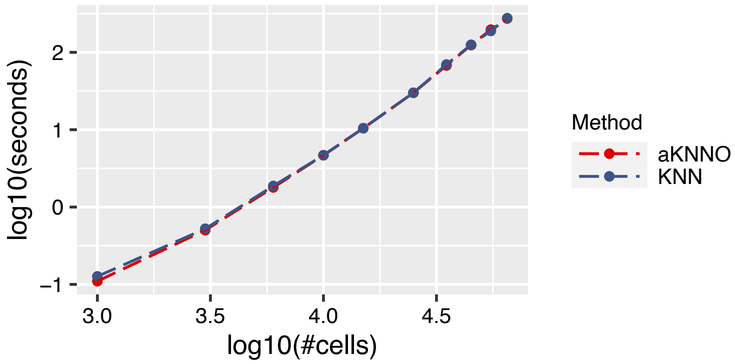

#### Application to single-cell RNA-seq from mouse intestinal

We analyzed one single-cell RNA-seq dataset from mouse intestine containing 559 cells[1]. aKNNO uncovered 10 clusters, which included all known cell types in the intestine epithelium (Supplementary Fig. S11a), including stem (cluster 3, n=65), TA (Transit Amplifying) (cluster 1, n=95), intermediate enterocytes (clusters 0 and 7), mature enterocytes (cluster 4, n=62), goblets (cluster 2, n=85), k cells (cluster 5, n=48), enteroendocrine (cluster 6, n=28), tuft (cluster 8,n=5), and Paneth cells (cluster 9, n=3). Their identities were verified by known markers, such as high level of *Ascl2* and *Lgr5* in stem, *Top2a* and *Mki67* in TA, *Fabp6* and *Apoa1* in mature enterocytes, *Muc2* and *Clca1* in goblets, *Gip* in k cells, *Chga* and *Chgb* in enteroendocrine, *Hck* and *Lrmp* in Tuft, and *Defa21* and *Defa22* in Paneth cells (Supplementary Fig. S11d). In comparison, KNN found five clusters and KNN_high detected nine clusters (Supplementary Figs. S11b and S11c). Neither KNN nor KNN_high identified those rare cell types successfully, including k cells, enteroendocrine, tuft, and paneth cells. They both failed to separate k cells and enteroendocrine (Supplementary Figs. S11b and S11c). In KNN and KNN_high, Paneth cells were hidden by goblets, while tuft cells were misclassified into enterocytes (Supplementary Figs. S11b and S11c).

GiniClust3 identified nine clusters in total, among of which six rare groups had less than 10 cells (Supplementary Fig. S11e). It failed to distinguish between intermediate and mature enterocytes, goblets and Paneth cells. It even misclassified stem, TA, k cells, enteroendocrine, and tuft cells into one cluster. The six rare clusters identified by GiniClust3 mixed with mature enterocytes and goblets cells in the UMAP embedding, suggesting they are not true rare (Supplementary Fig. S11e). GapClust obtained five rare clusters, two of which mapped to tuft and Paneth cells (Rare_2 and Rare_3 in the Supplementary Fig. S11f). The biggest rare cluster with 97 cells is goblet (Rare_5), which was not that rare.  FiRE quantified 25 cells as being rare (Supplementary Fig. S13g). It only found 1 out of 5 tuft cells and 15 out of 28 enteroendocrine to be rare. In summary, aKNNO is far more superior than KNN, KNN_high, GiniClust3, GapClust, and FiRE, to reveal all the known abundant and rare cell types in the intestinal epithelium (Supplementary Fig. S13h). aKNNO is very powerful in rare cells identification, which has the ability to detect the rare Paneth cell type with only three cells.

**Supplementary Fig. S11.** Application to single-cell RNAseq data from mouse intestine**.** The UMAP plot labeled by the manual annotation from aKNNO (a), KNN (b), KNN_high (c). (d) Dotplot of marker genes in clusters detected by aKNNO. (e) The UMAP plot labeled by the GiniClust3 result. (f) The UMAP plot labeled by the GapClust result. (g) The UMAP plot labeled by the FiRE result. (h) A summary of clusters identified by aKNNO, KNN, KNN_high, GiniClust3, GapClust, and FiRE.

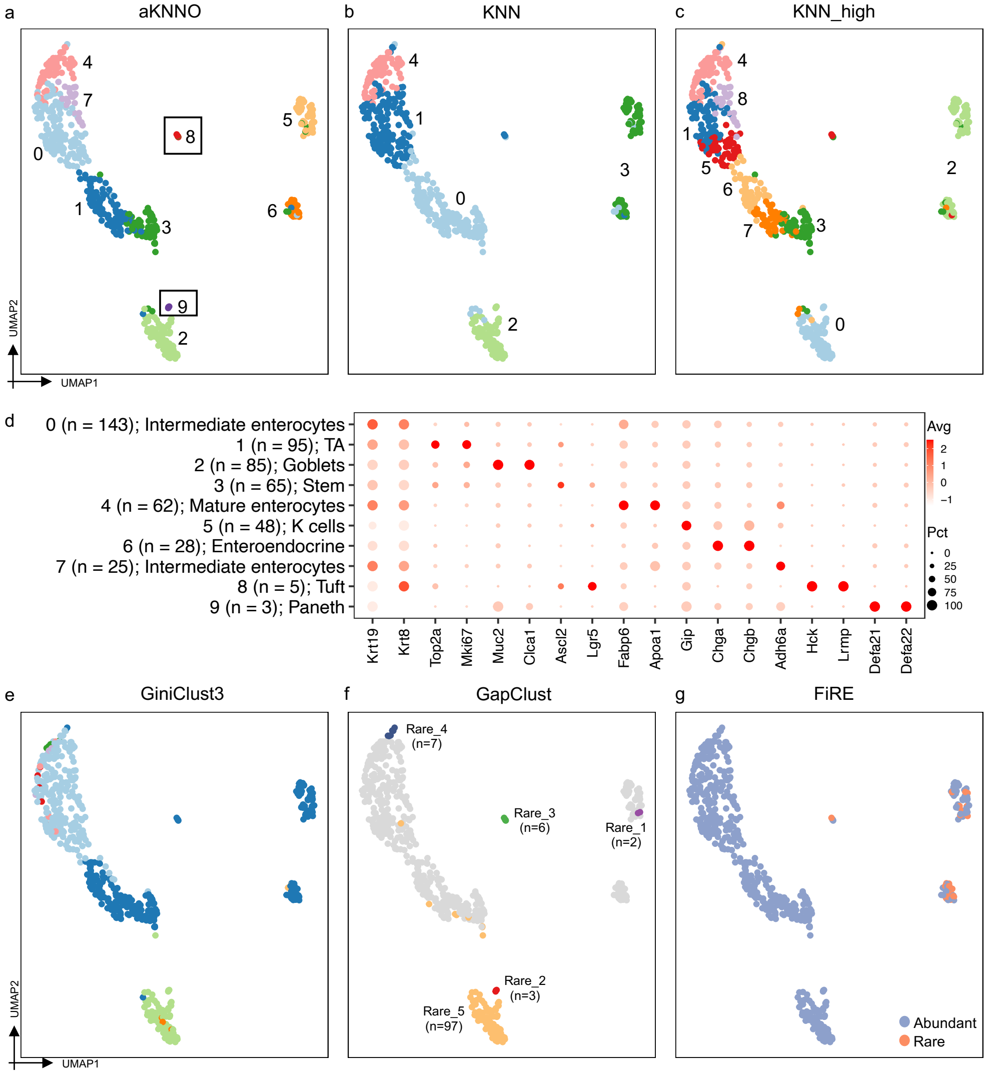


h

| aKNNO | KNN | KNN_high | GiniClust3 | GapClust | FiRE |
|---|---|---|---|---|---|
| 0 (n=143); Intermediate enterocyte | -- | -- | -- | -- | -- |
| 1 (n=95); TA | -- | -- | -- | -- | -- |
| 2 (n=85); Goblets | 2 (n=86) | 0 (n=86) | 2 (n=88) | Rare_5 (n=97) | -- |
| 3 (n=65); Stem | -- | 3 (n=67) | -- | -- | -- |
| 4 (n=62); Mature enterocyte | 4 (n=64) | 4 (n=63) | -- | -- | -- |
| 5 (n=48); K cells | -- | -- | -- | -- | -- |
| 6 (n=28); Enteroendocrine | -- | -- | -- | -- | -- |
| 7 (n=25); Intermediate enterocyte | -- | 8 (n=29) | -- | -- | -- |
| 8 (n=5); Tuft | -- | -- | -- | Rare_3 (n=6) | -- |
| 9 (n=3); Paneth | -- | -- | -- | Rare_2 (n=3) | -- |

**Application to single-cell RNA-seq from mouse habenula**

We analyzed single-cell RNA-seq derived from mouse habenula consisting of 1,149 cells, an epithalamic brain structure critical for processing and adapting to negative action outcomes[2]. aKNNO identified 17 clusters, while KNN and KNN_high found 11 and 13 groups, respectively (Supplementary Figs. S12a, S12b and S12c). aKNNO found more true rare cell types than KNN and KNN_high, including clusters 11 (n=23), 12 (n=17), 13 (n=15), 14 (n=10), 15 (n=2), and 16 (n=2). They were true cell types except the cluster 16, which could be validated by cell-type specific expression of known markers. Cluster 11 with high *Gap43* and *Slc17a6*, which are only expressed in LHb neurons[2]. Cluster 12 is myoepithelium cells with high expression of *Acta2* and *Myh11*. Cluster 13 is macrophages with high *Mrc1*. Cluster 14 is fibroblasts with specific expression of *Col1a1* and *Col3a1*. Cluster 15 with only two cells, it has the high expression of *Olig1* like other oligodendrocytes (clusters 4, 5, and 8), but with specific expression of *Top2a* and *Cdc20*, suggesting it is proliferating oligodendrocyte (Supplementary Fig. S12d). Cluster 16 shows much lower number of UMIs and genes compared to other clusters (Supplementary Fig. S12e), suggesting it is a cluster with empty droplets. KNN had similar clustering with aKNNO on abundant cells but missed all the six rare clusters (Supplementary Figs. S12a and S12b) (clusters 11-16 in the aKNNO). Although KNN_high identified much more clusters than KNN, it also failed to detect all the six rare clusters (Supplementary Fig. S12c).

GiniClust3 only identified five clusters in total, which misclassified distinct cell clusters into one group (Supplementary Fig. S12f). All the five clusters had greater than 100 cells, meaning no rare clusters were identified by GiniClust3. GapClust detected five rare clusters (Supplementary Fig. S12g). Three clusters were not that rare, among of which one having 123 cells from astrocytes (cluster 3 in the aKNNO), one with 126 cells from oligodendrocytes (cluster 4 in the aKNNO), and one with 33 clusters from microglia (cluster 9 in the aKNNO). Although GapClust identified fibroblasts (cluster 14 in the aKNNO) and doublets (cluster 16 in the aKNNO), it misclassified these two clusters into one (Rare_2 in the Supplementary Fig. S12g). FiRE found all cells to be common (Supplementary Fig. S12h). In summary, aKNNO is far more superior than KNN, KNN_high, GiniClust3, GapClust and FiRE in rare cells identification (Supplementary Fig. S12i).

**Supplementary Fig. S12.** Application to single-cell RNAseq data from mouse habenula**.** The UMAP plot labeled by the manual annotation from aKNNO (a), KNN (b), KNN_high (c). (d) Dotplot of genes marking the rare clusters detected by aKNNO. (e) The number of UMI and genes in the cluster 16 compared to other clusters. (f)The UMAP plot labeled by the GiniClust3 result. (g) The UMAP plot labeled by the GapClust result. (h) The UMAP plot labeled by the FiRE result. (i) A summary of rare clusters identified by aKNNO, KNN, KNN_high, GiniClust3, GapClust, and FiRE.

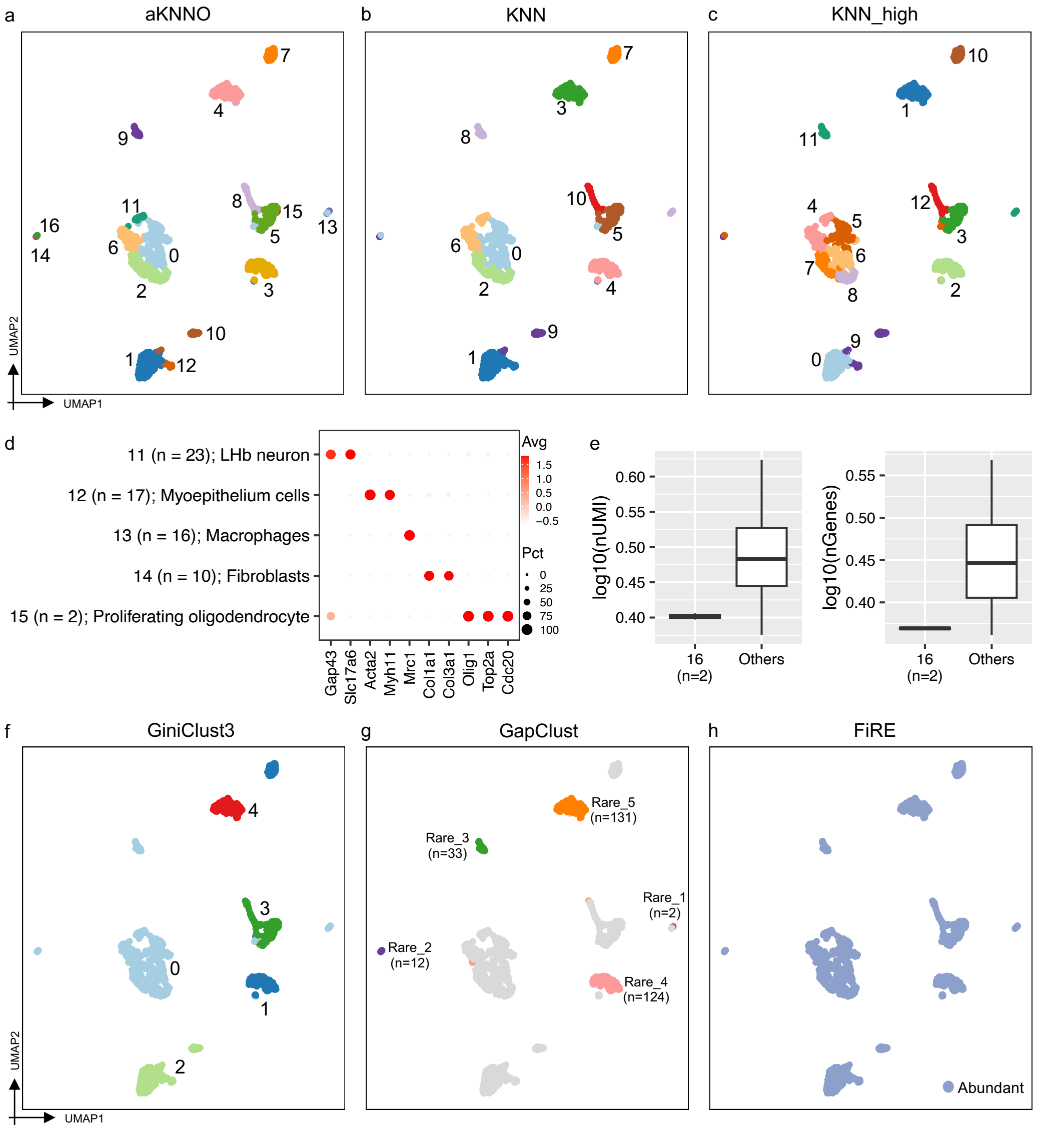


i

| aKNNO | KNN | KNN_high | GiniClust3 | GapClust | FiRE |
|---|---|---|---|---|---|
| 11 (n=23); LHb neuron | -- | -- | -- | -- | -- |
| 12 (n=17); Myoepithelium cells | -- | -- | -- | -- | -- |
| 13 (n=16); Macrophages | -- | -- | -- | -- | -- |
| 14 (n=10); Fibroblasts | -- | -- | -- | -- | -- |
| 15 (n=2); Proliferating oligodendrocyte | -- | -- | -- | -- | -- |
| 16 (n=2); Empty droplets | -- | -- | -- | -- | -- |

**Application to single-cell RNA-seq from intestinal organoids**

We applied aKNNO to one single-cell RNA-seq dataset from intestinal organoids, which were isolated and grown from mice intestinal crypts[3]. aKNNO and KNN_high both identified seven clusters from the 320 cells, while KNN only found four clusters (Supplementary Figs. S13a, S13b and S13c). The major difference between them is that aKNNO recognized two more rare clusters, cluster 5 (n=10) and cluster 6 (n=5). Cluster 5 had specific expression of *Neurog3* and *Chgb* (Supplementary Fig. S13d). This cluster has been reported to be enteroendocrine progenitor cells, where Neurog3 directs the program of enteroendocrine development[4,5]. Cluster 6 specifically expressed *Defa3* and *Defa5* (Supplementary Fig. S13d), which are known Paneth markers. In comparison, KNN failed to detect both two rare clusters, and KNN_high misclassified the two rare clusters into one group.

GiniClust3 identified four clusters (Supplementary Fig. S13f) and failed to detect rare enteroendocrine progenitor cells and Paneth cells. GapClust found two rare clusters, one with five cells and the other with two cells (Supplementary Fig. S13g). The one with five cells mapped to Paneth cells (Rare_2 in the Supplementary Fig. S13g). FiRE estimated nine cells to be rare (Supplementary Fig. S13h), but including only one of the five Paneth cells. The other eight cells mixed with abundant cells in the UMAP embedding, suggesting they were false. Among the two rare clusters identified by aKNNO (progenitor enteroendocrine and Paneth), only GapClust found Paneth cells, while other methods failed to detect both (Supplementary Fig. S13e).

**Supplementary Fig. S13.** Application to single-cell RNAseq data from intestinal organoids**.** The UMAP plot labeled by the manual annotation from aKNNO (a), KNN (b), KNN_high (c). (d) Dotplot of genes marking the two rare clusters. (e) A summary of two rare clusters identified by aKNNO, KNN, KNN_high, GiniClust3, GapClust, and FiRE. (f) The UMAP plot labeled by the GiniClust3 result. (g) The UMAP plot labeled by the GapClust result. (h) The UMAP plot labeled by the FiRE result.

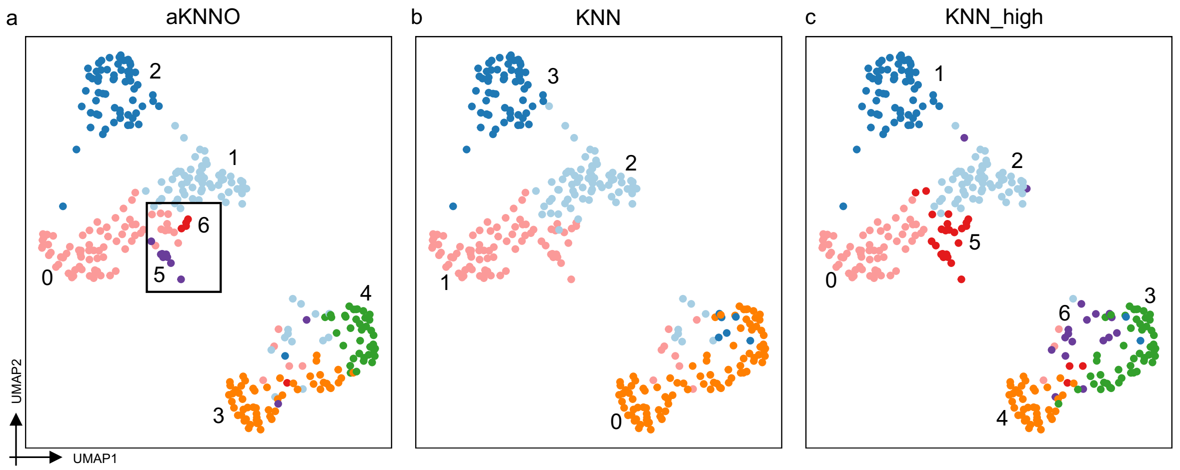


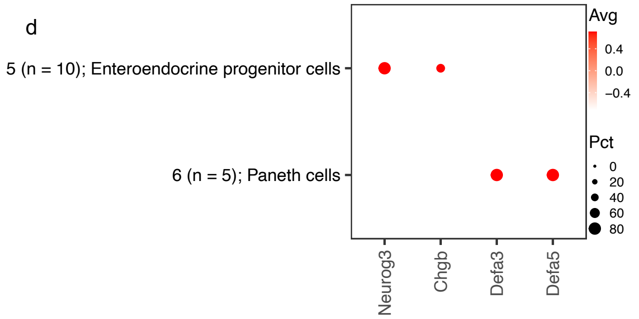


e

| aKNNO | KNN | KNN_high | GiniClust3 | GapClust | FiRE |
|---|---|---|---|---|---|
| 5 (n=10) | -- | -- | -- | -- | -- |
| 6 (n=5) | -- | -- | -- | Rare_2 (n=5) | -- |

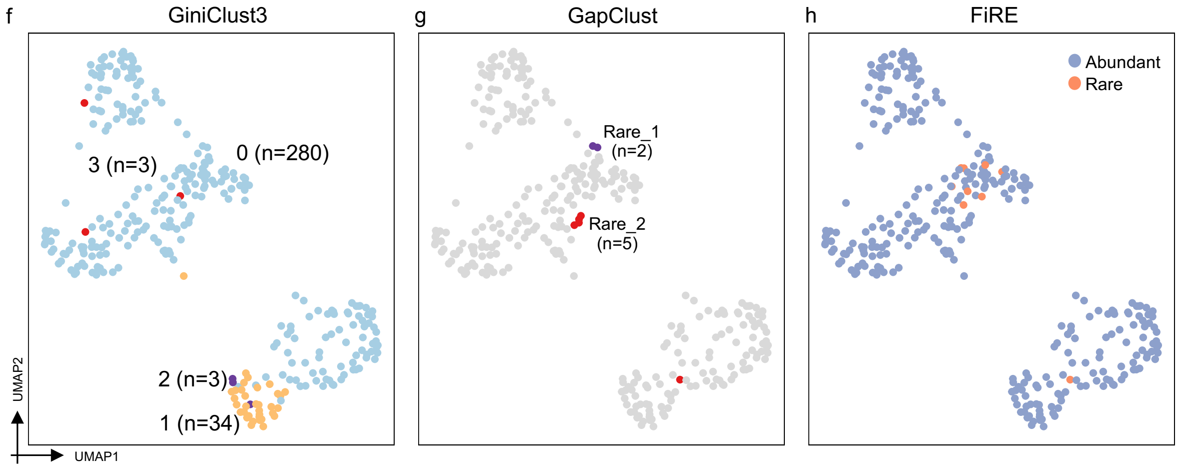

**Application to spatial transcriptomics from mouse coronal posterior brain**

We analyzed a 10x Visium dataset generated from mouse coronal posterior brain including 2,797 spots (Supplementary Fig. S14a). aKNNO, KNN and KNN_high detected 26, 16, and 27 clusters, respectively (Supplementary Figs. S14b, S14c and S14d). aKNNO and KNN_high both mapped dorsal hippocampus structures correctly, which involved four clusters corresponding to DG, CA3, CA1, and CA2_CA1. Their identities were supported by high expression of known markers (Supplementary Fig. S14e). KNN, however, only identified CA1 correctly but misclassified other regions into one cluster (Supplementary Figs. S14f, S14g and S14h).

The four rare clusters (clusters 22-25 in the Fig. S14i, n <30) all lined up with specific structures. Cluster 25 had high expression of *Tmem212*, *Ccdc153* and *Odf3b*, which has been reported as ependymal cells[6]. GiniClust3 only identified nine clusters in total, which lost fine anatomical structures of the mouse brain (Supplementary Fig. S14j). The four rare clusters (n<30) were scattered in the spatial region. GapClust found only two rare clusters (Rare_2, n=15; Rare_1, n=6) (Supplementary Fig. S14k). FiRE quantified 113 rare spots, among of which only eight spots mapped to the rare cluster 25 in the aKNNO and aligned to a specific anatomical pattern (Supplementary Fig. S14l). FiRE also quantified smooth muscle cells as being rare, which were not that rare compared to other clusters and also detected by aKNNO (cluster 19) (Supplementary Fig. S14m).

We compared aKNNO with the four integrative approaches, stLearn[7], SpaGCN[8], GraphST[9], and BayesSpace[10]. We set the number of their clusters to 26, matching the cluster number of aKNNO (Supplementary Figs. S15a-S15e). Although they all resolved anatomical structure of the mouse brain well, aKNNO outperformed the four integrative approaches in delineating fine-grained tissue structures. For example, aKNNO successfully identified four clusters (clusters 16, 22,23,24 in the Supplementary Fig. S15f), which stereotyped dorsal hippocampus structures DG, CA3, CA1, and CA2_CA1. BayesSpace detected four clusters, however, it showed an unclear structure of CA3 (cluster 2 in the Supplementary Fig. S15g). GraphST and stLearn only found two clusters, and SpaGCN recognized three clusters that were imprecise and merged into their surrounding regions (Supplementary Figs. S15h-S15j).

**Supplementary Fig. S14.** Application to 10x Visium spatial transcriptomics data from mouse coronal posterior brain. (a) H&E image. Spatial plot annotated by aKNNO (b), KNN (c) and KNN_high (d). (e) Dotplot of marker genes in the dorsal hippocampus structure. Spatial plot focusing on dorsal hippocampus structures annotated by aKNNO (f), KNN (g), and KNN_high (h). Spatial plot of clusters detected by GiniClust3 (i), GapClust (j), and FiRE (k). (l) A summary of rare clusters identified by aKNNO, KNN, KNN_high, GiniClust3, GapClust, and FiRE.

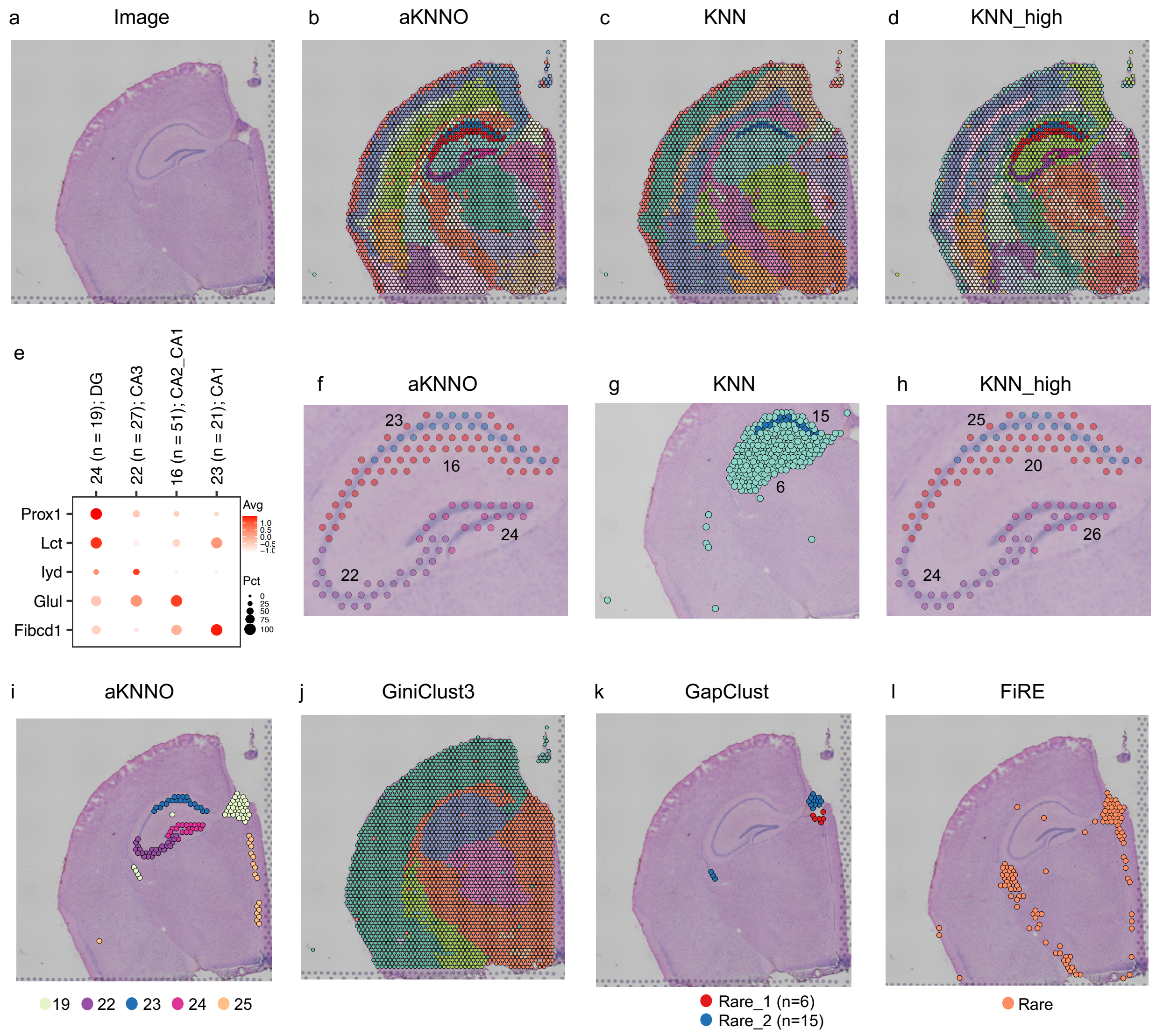


m

| Ref | aKNNO | KNN | KNN_high | GiniClust3 | GapClust | FiRE |
|---|---|---|---|---|---|---|
| -- | 19 (n=40); SMCs | 14 (n=33) | -- | -- | -- | Rare (n=35) |
| CA3_dorsal | 22 (n=27) | -- | 24 (n=25) | -- | -- | -- |
| CA1_dorsal | 23 (n=21) | 15 (n=20) | 25 (n=20) | -- | -- | -- |
| DG_dorsal | 24 (n=19) | -- | 26 (n=19) | -- | -- | -- |
| -- | 25 (n=18); Ependymal cells | -- | -- | -- | -- | Rare (n=10) |

**Supplementary Fig. S15.** Comparison between aKNNO, BayesSpace, GraphST, SpaGCN, and stLearn on the 10x Visium spatial transcriptomics data from mouse coronal posterior brain. Clustering of aKNNO (a), BayesSpace (b), GraphST (c), SpaGCN(d), and stLearn(e). Detailed view of clustering in the dorsal hippocampus structure in aKNNO (f), BayesSpace (g), GraphST (h), SpaGCN(i), and stLearn (j).

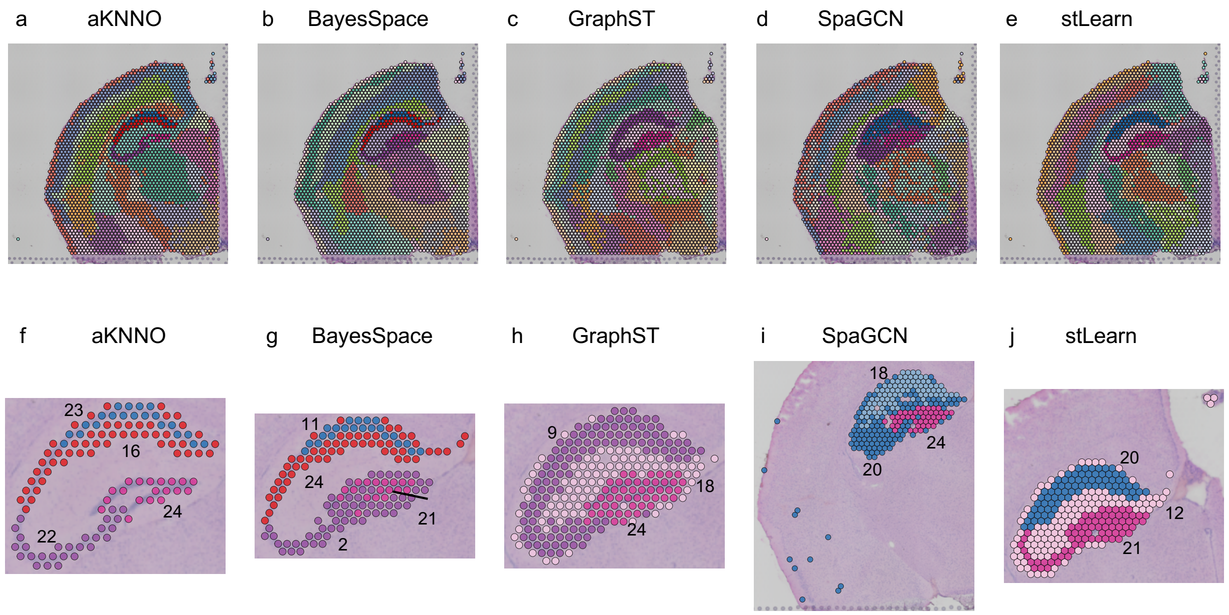

**Application to spatial transcriptomics from mouse main olfactory bulb**

We analyzed a 10x Visium dataset generated from mouse main olfactory bulb with 1,185 spots (Supplementary Fig. S16a). aKNNO, KNN and KNN_high detected 11, 8, and 13 clusters, respectively (Supplementary Fig. S16b, S16c and S16d). They all reconstructed seven-layer MOB structures well, including subependymal zone, granular cell layer, internal plexiform layer, mitral cell layer, external plexiform layer, glomerular layer, and outer nerve layer. In additional, aKNNO identified three additional clusters, one with specific expression of *Ptgds* (cluster 6, n=90) (Supplementary Fig. S16e and S16f), another with high level of *Hbb-bs*, *Hba-a1*, *Hba-a2*, and *Hbb-bt* (cluster 8, n=52) (Supplementary Fig. S16e and S16f), and the third one with specific expression of *Tpbg* (cluster 9, n=31). Cluster 6 was olfactory sheathing cells, cluster 8 was located in the glomerular layer and mapped to glomerular capillaries[11], and cluster 9 has been previously reported as 5T4 granule cells[12]. In comparison, KNN missed all the three clusters (Supplementary Fig. S16g), while KNN_high failed to detect olfactory sheathing cells (Supplementary Fig. S16h).

GiniClust3 identified five clusters, which lost the seven-layer MOB structures (Supplementary Fig. S16i). FiRE found 17 rare cells (Supplementary Fig. S16j), all of which scattered in the mob region and didn't map to any anatomical cell types. GapClust didn't run successfully on this dataset. In summary, methods except aKNNO and KNN_high failed to detect any rare clusters (Supplementary Fig. S16k).

We compared aKNNO with the four integrative approaches, stLearn[7], SpaGCN[8], GraphST[9], and BayesSpace[10]. We set the number of their clusters to 11, matching the cluster number of aKNNO (Supplementary Figs. S17a-S17e). aKNNO, BayesSpace, and SpaGCN defined the MOB seven-layer structure well, while GraphST and stLearn failed to identify subependymal zone.

**Supplementary Fig. S16.** Application to 10x Visium spatial transcriptomics data from mouse main olfactory bulb. (a) H&E image. Spatial plot annotated by aKNNO (b), KNN (c) and KNN_high (d). (e) Dotplot of marker genes in three rare clusters. Spatial plot focusing on the three rare clusters annotated by aKNNO (f), KNN (g), and KNN_high (h). Spatial plot of clusters detected by GiniClust3 (i), and FiRE (j). (k) A summary of rare clusters identified by aKNNO, KNN, KNN_high, GiniClust3, and FiRE.

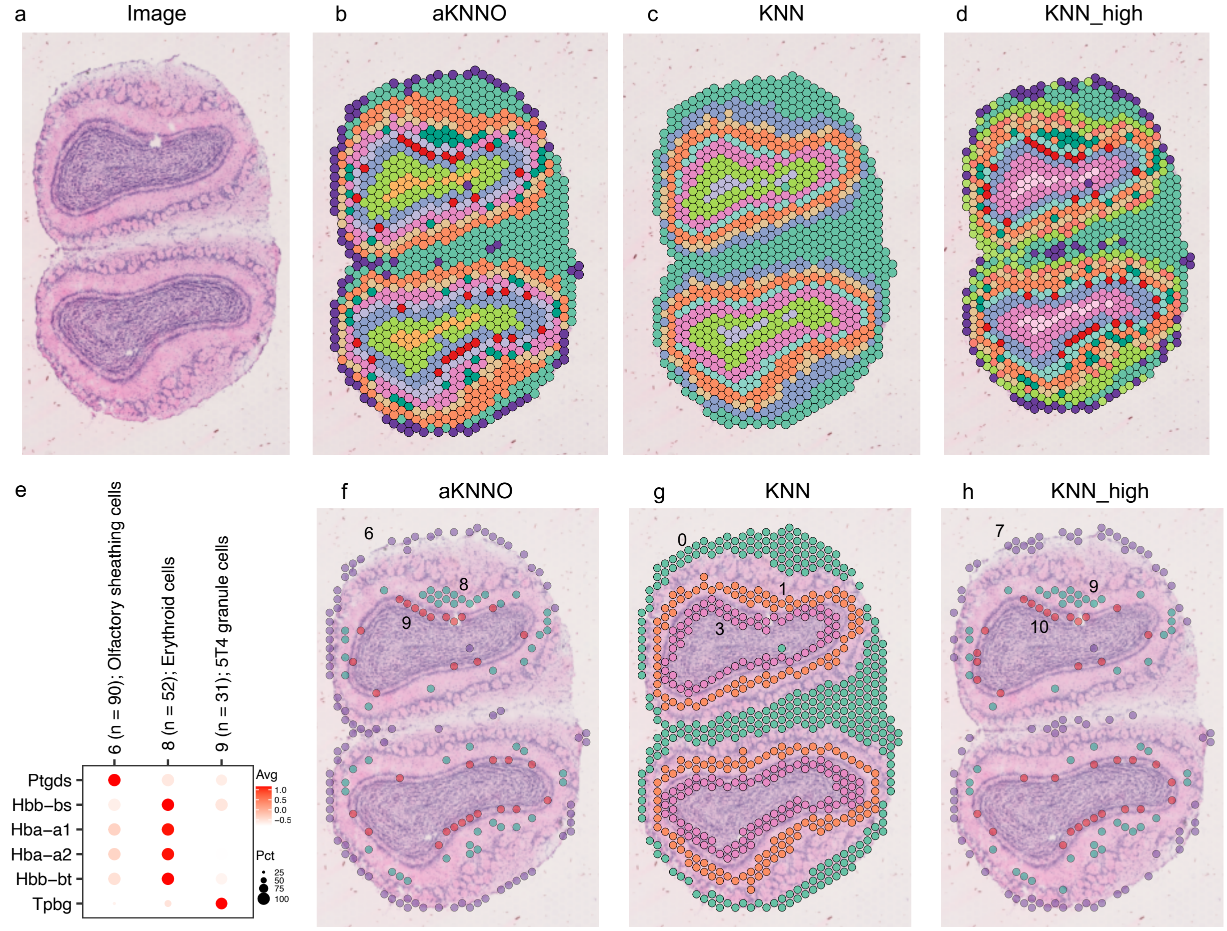


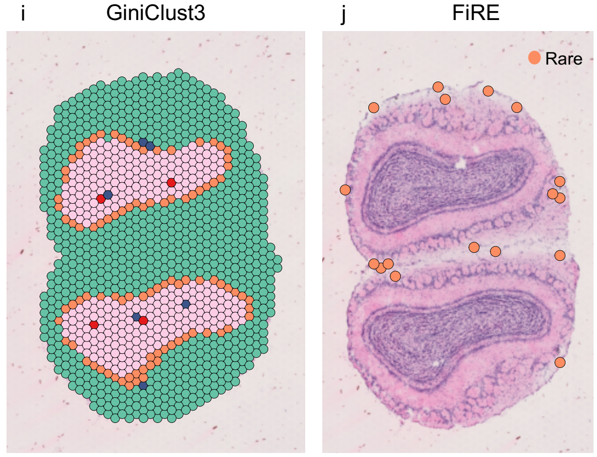


k

| aKNNO | KNN | KNN_high | GiniClust3 | FiRE |
|---|---|---|---|---|
| 6 (n=90) | -- | 7 (n=80) | -- | -- |
| 8 (n=52) | -- | 9 (n=50) | -- | -- |
| 9 (n=31) | -- | 10 (n=35) | -- | -- |

**Supplementary Fig. S17.** Comparison between aKNNO, BayesSpace, GraphST, SpaGCN, and stLearn on the 10x Visium spatial transcriptomics data from main olfactory bulb. Clustering of aKNNO (a), BayesSpace (b), GraphST (c), SpaGCN(d), and stLearn(e).

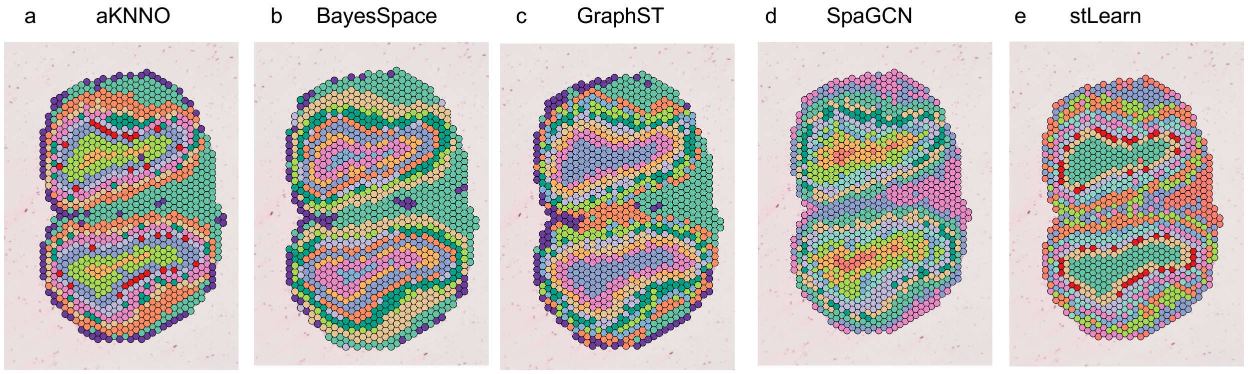

**Identification of rare lymphatic endothelial cells from 12 pan-cancer datasets**

Endothelial cells (ECs) have been proved to promote tumor angiogenesis[13], also evolve in immune regulation in the tumor microenvironment[14]. Based on previous pan-cancer scRNAseq data analysis, ECs in tumor were classified into 5 subtypes, including ESM1 tip cells only resided in malignant tissue, ACKR1 high venous ECs enriched in tumor, CA4 capillary ECs enriched in normal tissue, FBLN5 arterial ECs and PROX1 lymphatic ECs (LECs)[15,16]. Specifically, LECs facilitate cancer invasion in the tumor lymphatic metastasis, in which malignant cells need to squeeze between LEC junctions and move along LECs towards distant organs[17].

We applied aKNNO to identify LECs in 12 datasets across 11 cancer types[16,18-26]. Although LECs were rare, aKNNO reported LECs in all datasets (Supplementary Fig. S18a). The number of LECs ranged from 16 to 158, with the percentage even less than 0.15%. For example, in the breast cancer dataset (BRCA_GSE148673), 16 out of 10,359 cells were identified to be LEC (0.15%). In the non-small lung cancer (NSCLC_EMTAB61469), 80 out of 40,218 cells were recognized as LEC (0.2%) (Supplementary Fig. S18a). The detailed view of clustering for each dataset was shown in Supplementary Figs. S19 and S20). For example, aKNNO identified two endothelial clusters in the BRCA_GSE148673 dataset, which were cluster 23 ESM1 tip cells with 107 cells and cluster 40 LECs with 16 cells (Supplementary Fig. S19a). Consistently, *ESM1* and *NID2* were highly expressed in the cluster 23, while *PDPN* and *PROX1* were highly expressed in the cluster 40 (Supplementary Fig. S19a). We estimated the percentage of LECs in the endothelial cells across cancer types (Supplementary Fig. S18b). Melanoma had the highest percentage of LECs, followed by UVM and HNSC, which is consistent with one previous study on pan-cancer LECs proportions [27].

**Supplementary Fig. S18.** Application to 12 single-cell RNAseq data from 11 cancer types. (a) A summary of each dataset and aKNNO results. (b) Percentage of LECs in the endothelial cells across the 12 datasets.

a

| Dataset | Total cell number | Total aKNNO clusters | LECs | Percentage |
|---|---|---|---|---|
| BRCA_GSE148673 | 10,359 | 44 | 40 (n=16) | 0.00154 |
| CHOL_GSE142784 | 10,526 | 27 | 15 (n=122) | 0.0116 |
| CRC_EMTAB8107 | 34,284 | 37 | 36 (n=158) | 0.0046 |
| ESCA_GSE173950 | 9,544 | 36 | 31 (n=46) | 0.00482 |
| HNSC_GSE103322 | 5,902 | 40 | 31 (n=46) | 0.0078 |
| Melanoma_GSE72056 | 4,645 | 28 | 21 (n=41) | 0.00883 |
| NHL_GSE128531 | 30,497 | 48 | 45 (n=90) | 0.00295 |
| NHL_GSE147944 | 11,245 | 34 | 31 (n=41) | 0.00365 |
| NSCLC_EMTAB6149 | 40,218 | 43 | 41 (n=80) | 0.00199 |
| OV_EMTAB8107 | 32,386 | 46 | 37 (n=129) | 0.00371 |
| STAD_GSE167297 | 22,464 | 34 | 33 (n=24) | 0.00107 |
| UVM_GSE138433 | 12,682 | 29 | 28 (n=18) | 0.00142 |

b

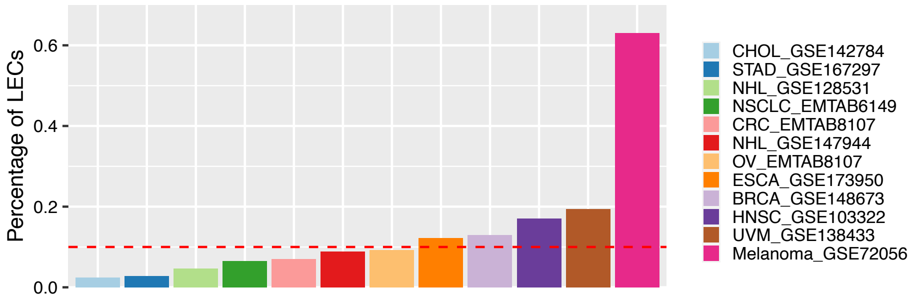

**Supplementary Fig. S19.** UMAP plots annotated by aKNNO results and expression of *PDPN* and *PROX1* marking LECs in BRCA(a), CHOL(b), CRC(c), ESCA(d), HNSC (e) and melanoma (f).

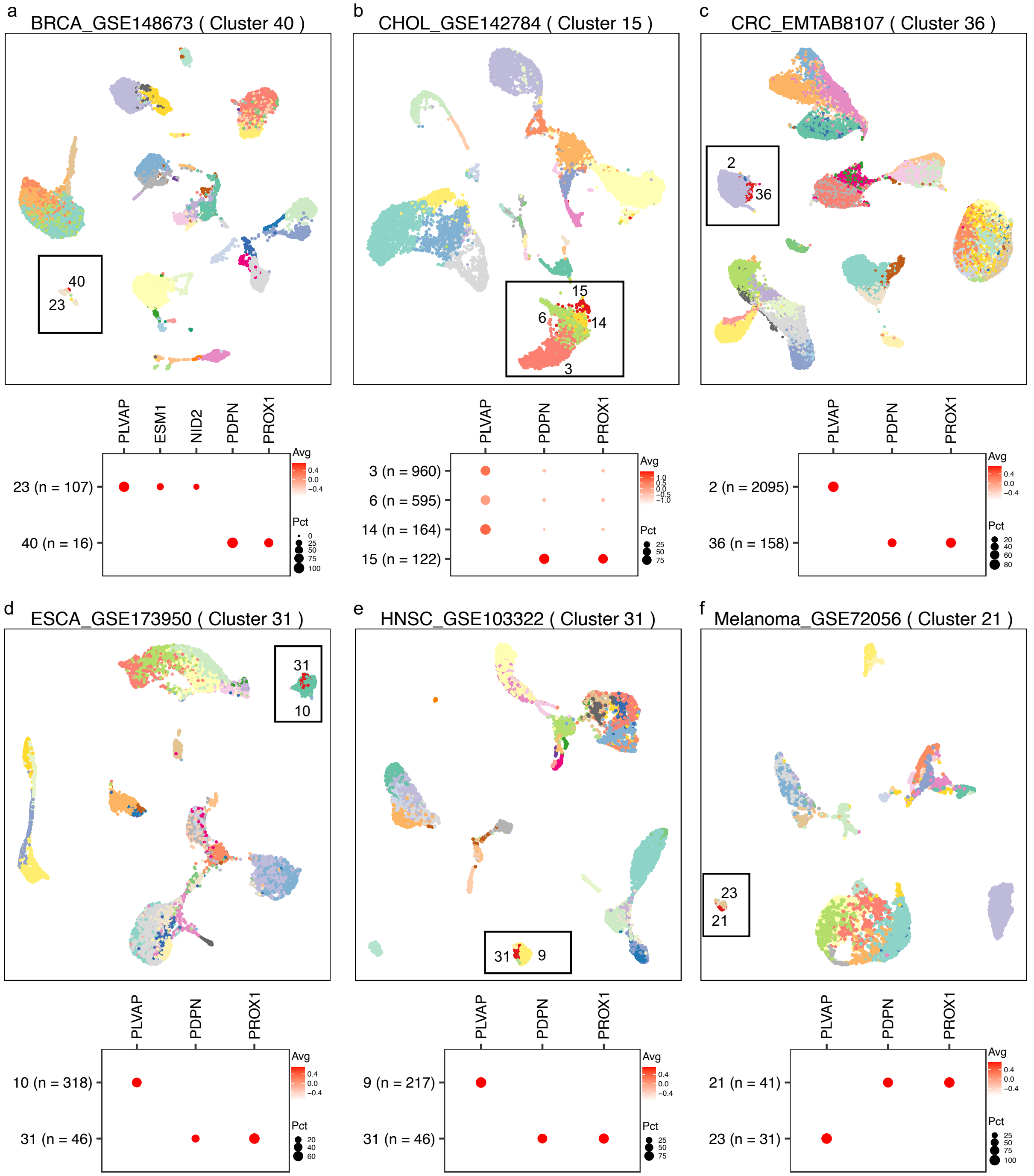

**Supplementary Fig. S20.** UMAP plots annotated by aKNNO results and expression of *PDPN* and *PROX1* marking LECs in NHL_GSE128531(a), NHL_GSE147944(b), NSCLC(c), OV(d), STAD (e) and UVM (f).

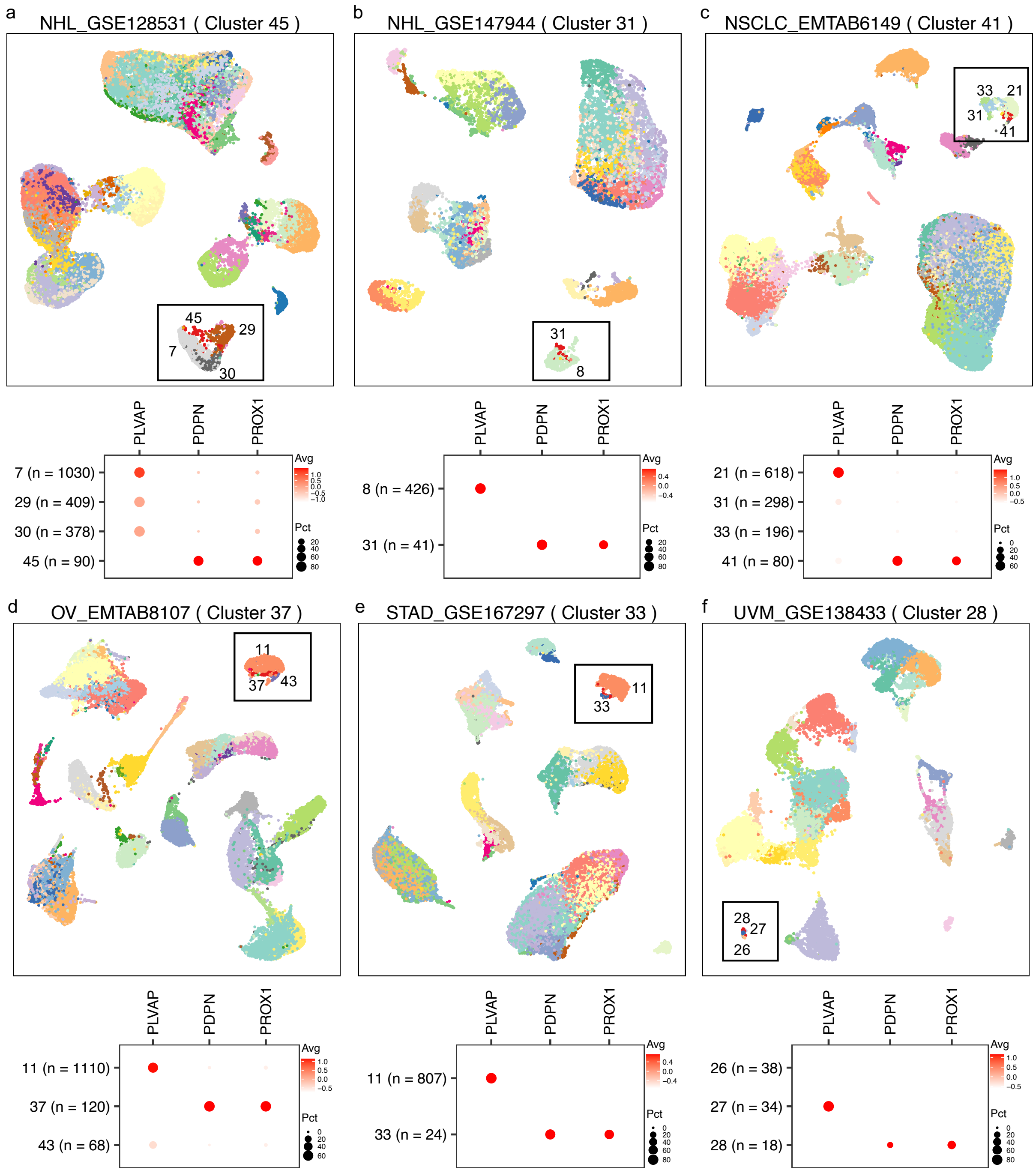
